# Female-biased sex ratios despite stable genetic sex determination across a climatic gradient in a marine fish

**DOI:** 10.64898/2026.02.16.706123

**Authors:** S. Lorena Ament-Velásquez, Trond Amundsen, Sebastian Wacker, Linn A.S. Aresvik, Halvor Knutsen, Marte Sodeland, Ignas Bunikis, Elisabet Forsgren, Anne Christine Utne-Palm, Qiaowei Pan, Ivain Martinossi-Allibert

**Affiliations:** Department of Zoology, Stockholm University, 106 91 Stockholm, Sweden; Department of Biology, Norwegian University of Science and Technology (NTNU), Høgskoleringen 5, 7034 Trondheim, Norway; Norwegian Institute for Nature Research (NINA), Høgskoleringen 9, 7034 Trondheim, Norway; Institute of Marine Research, Nye Flødevigveien 20, 4817 His, Norway; Centre for Coastal Research, Department of Natural Sciences, University of Agder, 4630 Kristiansand, Norway; Department of Immunology, Genetics and Pathology, Uppsala Genome Center, National Genomics Infrastructure Hosted by SciLifeLab, Uppsala University, Uppsala, Sweden; Fish Capture Division, Institute of Marine Research (IMR), Nordnesgaten 50, 5005 Bergen, Norway; Max Planck Institute for Biology, 72076, Tübingen, Germany; Faculty of Biosciences and Aquaculture, Nord University, Bodø, Norway

**Keywords:** Adult sex ratio, coastal ecosystem, sex determination, primary sex ratio, latitude

## Abstract

Adult sex ratio (ASR) is a fundamental parameter shaping population dynamics and evolutionary trajectories. In theory, ASR is governed by the sex determination (SD) system and sex-specific mortality. Yet, ASR is often hard to predict in natural populations because of ecological effects on its determinants. We address this ecological complexity by investigating the drivers of ASR across a steep climatic gradient in Scandinavian populations of the two-spotted goby *Pomatoschistus flavescens*.

Demographic surveys accounting for over 25000 fish in 30 locations, spanning 10° of latitude, revealed a general female-bias of the ASR, *ca.* 75%. To assess the contribution of SD, we combined reduced representation sequencing of 180 adults with our novel long-read male genome assembly. We characterised the genetic architecture of an XX/XY SD system, and identified a male-specific duplicate from the TGF-β signaling pathway, *amhr2y,* as a candidate sex-determining gene. Subsequent genotyping of eggs and juveniles showed consistently unbiased sex ratios across latitudes, implying that the female-biased arises at later life stages. Interpreted in the light of historical data, our findings allow us to reject environmental sex determination and sex-biased mortality as important drivers of ASR bias in this species. We propose instead that the female bias originates from sex-specific patterns of habitat distribution, which could result from ecological conditions interacting with the distinct reproductive strategies of females and males. Our results underscore the need to go beyond primary theoretical expectations to understand fundamental demographic parameters within their relevant ecological context.

## Introduction

Variation in sex ratio can have profound consequences for both demographic and evolutionary processes. Sexual selection, a potent source of evolutionary innovation (Andersson 1994), is particularly sensitive to fluctuations in the sex ratio of reproductively active individuals (Operational Sex Ratio, OSR, Emlen and Oring 1977, Kappeler et al. 2023). More broadly, the Adult Sex Ratio (ASR) can influence life-history strategies, mating competition tactics, and parental care on both ecological and evolutionary timescales (Székely et al. 2014, Eberhart-Phillips et al. 2018, Kappeler et al. 2023). ASR is also a key demographic variable that can affect both population productivity and effective population size, and as such must be accounted for in conservation strategies (Kappeler et al. 2023). Despite its importance, the drivers of ASR variation in natural populations remain poorly understood (Székely et al. 2014).

Over the last 20 years, the two-spotted goby *Pomatoschistus flavescens* has been an instrumental model system for the study of how sex ratios affect sexual selection in wild populations (Forsgren et al. 2004, Myhre et al. 2012, Wacker et al. 2014, Utne-Palm et al. 2015, Martinossi-Allibert et al. 2025a). *Pomatoschistus flavescens* is a very abundant planktivorous fish, gathering in large numbers near the shore during breeding, which makes it both an important component of coastal ecosystems in the North-East Atlantic (Fosså 1991) and a tractable field system (Amundsen 2018). This small fish with male-only parental care features colourful ornaments in both sexes (Amundsen 2018), which are involved in complex courtship and competition behaviours responding to fluctuations in the local sex ratio of breeders (Forsgren et al. 2004, Martinossi-Allibert et al. 2025a).

Strikingly, heavily female-biased ASR, *circa* 75% females, have been observed in the historical study population of Western Sweden (Lat. 58°N, Forsgren et al. 2004), but it remains unclear whether this ASR skew is representative of *P. flavescens* populations, and what might be causing it. The study of *P. flavescens* populations has recently been expanded to the northern limit of the species range (Lat. 68°N), revealing important shifts in mating behavior and reproductive strategy along the steep climatic gradient of the Scandinavian coastline (Martinossi-Allibert et al. 2025a and b). Exploring ASR along this gradient opens a unique opportunity to disentangle the contributions of genetic architecture, life-history, and ecological variables into the determination of ASR in wild populations of this nest-brooding marine fish.

Differences in mortality rates between the sexes, from birth to adulthood and during the adult period, are perhaps the most common cause of ASR bias (Székely et al. 2014). Sex-specific mortality results from the distinct reproductive strategies of females and males, implying differences in growth trajectories, predation risks (Magnhagen 1991), dispersal propensity (Trochet et al. 2016), and levels of parental investment (Arnqvist and Rowe 2005). This is particularly well studied in birds, where Donald (2007) reported 65% of published estimates giving biased ASR and an average excess of males of 33%, driven by overall higher female mortality. Independently of sex-specific mortality, biased sex ratio at birth can also contribute to biased ASR. Classical evolutionary theory predicts that, assuming equal cost of producing males and females, we should expect a balanced sex ratio at fertilisation (Düsing 1884, Edwards 2000, Seger and Stubblefield 2002). This is the case in most species with Genetic Sex Determination (GSD), but in species with environmental sex determination, distortion of the primary sex ratio is frequent, following fluctuations of the environmental cue.

Even though the majority of fish studied with separate sexes have GSD (Bachtrog et al. 2014), environmental parameters such as temperature frequently interfere with sex determination (Ospina-Alvarez and Piferrer 2008, Geffroy and Wedekind 2020, Lema et al. 2024). Consequently, the role of genetic versus environmental factors in fish sex determination can be hard to establish without an overview of the entire species range and its associated climatic conditions. For instance, Duffy et al. (2015) reported that in populations of Atlantic silverside *Menidia menidia* along the East coast of North America, the degree of environmental contribution to sex determination varied across the latitudinal range. In the Southern flounder *Paralichthys lethostigma*, the XX/XY sex determination system is affected by low and high temperature extremes in the natural species range, leading to masculinization of XX individuals (Honeycutt et al. 2019).

Disentangling the determinants of ASR in fish can thus be challenging, since sex determination system and sex-specific mortality can both be sensitive to ecological conditions. To understand ASR in natural populations, it is thus necessary to (i) know the sex determination system, and (ii) have an overview of the sex ratio of early life stages, as well as (iii) the ASR itself, across a range of environmental conditions. Starting from the previously published observations of heavily female-biased ASR in *P. flavescens*, we include these three aspects in a comprehensive investigation of ASR patterns across six populations covering the steep climatic gradient of the Scandinavian range of this species (Lat. 58°N to Lat. 68°N). To elucidate the sex determination system of *P. flavescens*, we generated a reference genome assembly from a male, as well as short-read data from four individuals of known sex, and combined them with population genomics data from 178 adults. In parallel, we conducted a large-scale field study in 2022, to get an overview of sex ratios at different life stages across the latitudinal gradient of the Scandinavian range. Finally, we put the 2022 ASR data in context with field data collected over the past two decades in different locations in the North Sea and the Baltic Sea, and using a variety of methods.

Together, these converging lines of evidence provide a comprehensive insight into sex ratio fluctuations in populations of a widespread marine fish of the Northeast Atlantic. Our study highlights the complexity of ASR dynamics in natural settings, which has cascading implications for predictive demography and evolutionary dynamics.

## Methods

### 1. Field sampling

#### 1.1 Demographic surveys in 2022

From April 24th 2022 to July 25th 2022, population censuses were conducted in the six study populations (**Figure 1a**, triangles). Each population was replicated by five study transects located within a few kilometers of each other. Each of the 30 transects was sampled three times, representing early season, peak season, and late season samplings. Transects were on average 120 m long, and established along the shore in the breeding habitat (kelp and other seaweed). The dates of sampling and exact locations of the study transects are given in Supplementary File F1. First, observers swam along the transect and counted the number of fish of each sex in a corridor of approximately 1 m around the observer. This was done twice on each sampling occurrence. In total, 25771 fish were recorded. Then, fish were caught individually or in small groups by snorkelling field workers using dip nets, and brought back to the shore for sex identification (In total 8374 fish caught). To minimize bias in the catch as much as possible, the protocol instructed the snorkeller to swim out of the breeding habitat, then back in and target the first fish that was observed. Phenotypic sexing was done based on secondary sexual characteristics: males were identified based on extended and darkened dorsal and anal fins, and females identified based on the absence of fin pigmentation, and in many cases the roundness of the belly associated with egg maturation. Most of the adult fish caught could be easily sexed (97% of captured fish), and the ones that could not be sexed, sexually immature adults, belonged to the two northernmost populations where postponement of reproduction is frequent (Martinossi-Allibert et al. 2025b).

**Figure 1.**
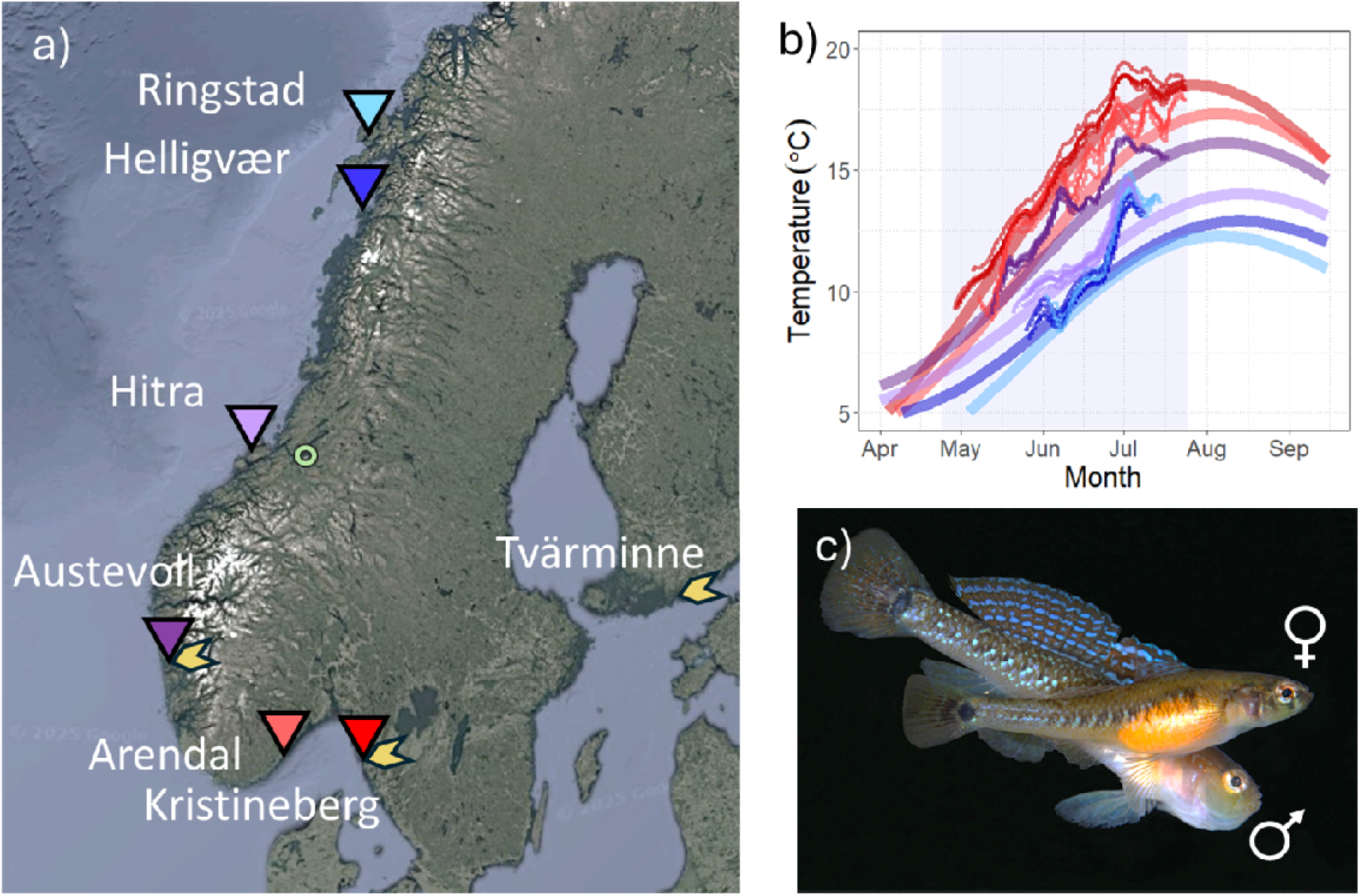
Location of the study populations (a), temperature in the breeding habitat (b), and Pomatoschistus flavescens pair in courtship (c). In (a) the location and names of the study populations are given on a map of Norway (background map from Google Maps 2025). The study populations for the 2022 campaign are shown with standing triangles, the three historical populations with leaning yellow arrowheads, and the sampling location of the male collected for genome assembly by a green ring, in the Trondheim fjord. In (b), the temperature in the breeding habitat is given for the 2022 populations, with the colors matching symbols in (a). The thin lines represent sliding averages of the temperature recorded hourly in the field over the sampling period (highlighted in blue). The thick lines show a sinusoidal fit to 17 years of sea surface temperature from the Norkyst model for these locations (2005-2022). In (c) a female (foreground) and male in courtship are shown (Photograph by Nils Aukan). Males are identified by fin morphology and coloration, and females either by belly shape and coloration if eggs are present, or by absence of fin coloration otherwise.

#### 1.2 Collection of genomics samples in 2022

During the same field effort as for the demographic survey, one fish from each catch and each sex was euthanised with a lethal dose of MS222 and preserved in 95% ethanol. The rest of the catch was released on site. In total, this represented a balanced sample of 180 fish (6 populations times 5 study sites times 3 sampling times 2 sexes). These samples were used for ddRAD sequencing, with the primary goal of assessing population structure across the range (see Martinossi-Allibert et al. 2025b). In addition, 100 adults from the northernmost population of Ringstad that could not be phenotypically sexed due to the lack of secondary sexual characteristics were euthanised in excess MS222 and preserved in 95% ethanol, in order to allow for subsequent sex attribution by genetic markers. These samples were sexed using genotype by sequencing. Furthermore, in three of the five study sites of each population, artificial nests were set up during early season sampling within the natural populations, and eggs were collected in these nests during the peak and late season samplings. Eggs and occasionally larvae, if the eggs hatched during collection, were preserved in 95% ethanol. They were then genotyped for sex using genotype by sequencing.

#### 1.3 Earlier field surveys (2007-2019)

##### Observation transect in Sweden, Norway and Finland

Population censuses were conducted by snorkeling transects along the coastline in prime breeding habitat at three localities; Kristineberg Marine Research Station, Sweden (2007, 2008, 2010), Espegrend Marine Biological Station, Bergen, Norway (2007, 2008), and Tvärminne, Finland (2008). In each population, 3 study sites were sampled (1 transect per site in 2007, and 2 transects per site later on) at 2 to 3 occurrences over the breeding season, by two observers each time. Transect length ranged from 36m to 172m. Adult fish observed within 0,5 m to the sides and 1 m below the observer were counted and sexed as the observer slowly swam along the transect. For more details, see Forsgren et al. (2004).

##### Beach seine samplings in Bergen

Fish were sampled during early (May) and late (July) breeding using a 30 m × 3 m beach seine with 3–5 mm square mesh. The sein was modified with additional weights on the bottom line and floats on the top line to ensure it followed the seabed through macroalgae, thereby providing a representative sample of nest-holding males occupying nests on the bottom and among macroalgae. Sampling took place at three west-coast sites north of Bergen: inner Fanafjord, outer Fanafjord, and Kvalen in Raunefjord.

### 2. Genomics

#### 2.1 Whole genome sequencing and assembly

##### Illumina sequencing of two individuals of each sex

Four adult individuals collected during the 2022 field season were chosen for short-read sequencing. The fish were euthanised in the field with excess MS222 and preserved in 95% ethanol at room temperature until 2024. To ensure that the phenotypic sex was correct, we selected males that were captured while guarding nests (females do not contribute to parental care) and females that were carrying visible eggs. One individual of each sex originated from a northern population (Helligvær) and one of each sex from the southernmost population of the sampling effort (Kristineberg).

DNA was extracted from muscle and fin tissue of the tail, using the MagAttract HMW DNA Kit (Qiagen) and further purified with the Genomic DNA Clean & Concentrator Kit (Zymo Research) to remove excessive pigment. The resulting DNA was processed into 150 bp paired-end Illumina DNA PCR-free libraries and sequenced with a NovaSeqX Plus system at the National Genomics Infrastructure (NGI) in Stockholm, Sweden.

We removed duplicates from the raw Illumina reads using fastp v. 0.24.0 (Chen 2023) with parameters --disable_quality_filtering --disable_adapter_trimming --disable_trim_poly_g --dedup --dup_calc_accuracy 4. The output was re-run again on fastp to perform general quality control, with parameters --length_required 80 --cut_front --cut_tail -f 20. These were considered “clean” Illumina reads.

##### Pacific Biosciences (PacBio) sequencing of one male

One individual male, measuring 57 mm and weighing 1.68 g, was captured in the bay of Korsvika, in the Trondheim fjord near Trondheim, Norway (Lat. 63.449650 N, 10.431350 E, **Figure 1a**) on the 23rd of September 2024 and labeled as TH1. The fish was euthanised with excess MS222 and approximately 200 mg of tail tissue was cut and frozen in liquid nitrogen, and subsequently preserved at -80°C or on dry ice.

High-molecular-weight (HMW) DNA was extracted from frozen tail muscle using the Nanobind PanDNA kit (103-260-000, PacBio®) following “Extracting DNA from animal tissue using the Nanobind®PanDNA kit - Procedure & checklist” (102-574-600, REV04 AUG2024). A total of ∼200 mg of tissue was homogenized in 1.5 ml of CT buffer, divided into 3 tubes with 750, 500, and 250 µl, respectively, and brought up with CT buffer to a total volume of 750 µl if needed. For each reaction, DNA was eluted in 200 µl kit EB and left at room temperature with gentle mixing at 100 rpm on a platform rocker for several days to increase homogeneity before performing quality control (QC). The best extraction was processed into a SMRTbell library at the Uppsala Genome Center, Sweden. Sequencing was performed on two Revio SMRT cells of the Pacific Biosciences (PacBio) Revio technology platform. Resulting HiFi reads were assembled into Hifiasm v. 0.24.0 (Chen et al. 2022) with parameters --dual-scaf --telo-m CCCTAA. The mitochondrial contig was taken from the Hifiasm primary assembly and circularized using MitoHiFi v. 3.2.2 (Uliano-Silva et al. 2023) with parameters -f NC_052761.1.fasta -g NC_052761.1.gb.

The Hifiasm ouput contains three assemblies: the primary assembly (with collapsed haplotypes) and two haplotype-resolved assemblies. For downstream analyses, we cleaned the primary assembly by replacing any mitochondrial contigs with the MitoHiFi output. We further removed all contigs that had no read mapping from the four Illumina samples, indicating that they correspond to contamination rather than true goby DNA. To this end, the cleaned Illumina reads were mapped using BWA v. 0.7.18 (Li and Durbin 2009) and piped to SAMtools v. 1.21 (Danecek et al. 2021) to produce sorted BAM files. We applied Picard v. 3.3.0 (http://broadinstitute.github.io/picard/) with the optical duplicate pixel distance set to 2500 to mark duplicates. We used Cramino v. 1.0.0 (De Coster and Rademakers 2023) to obtain mean depth of coverage estimates from the BAM files. This strategy removed 212 small contigs. Based on the statistics of BUSCO v. 5.8.2 (Manni et al. 2021) and the actinopterygii_odb12 database (n:7207), the final TH1 primary assembly was comparable (C:95.6%[S:93.5%,D:2.1%],F:1.4%,M:3.0%) to the initial primary assembly (C:95.6%[S:93.8%,D:1.7%],F:1.4%,M:3.0%), and to the reference genome (C:95.5%[S:95.0%,D:0.5%],F:1.3%,M:3.2%) of a French specimen (accession number OZ251433.1), here after fGobFla1. The final TH1 primary assembly contains no gaps. However the scaffold h1tg000060l, corresponding to the Y haplotype in the haplotype 1 assembly, contains one gap located 500 Kb upstream of the SD region.

To verify that fGobFla1 and TH1 belong to the same species, we extracted their *cox1* mitochondrial gene, along with the homolog from a published mitogenome from a Danish individual (accession number NC_052761.1) and closely related goby species. To this end, we used the *cox1* sequence from the Danish *P. flavescens* as a query to extract the homologs from our assembled mitogenome, the mitochondrial contig of fGobFla1, and the whole-genome assembly of *P. minutus* (Leder et al. 2021). Additionally, we performed BLASTn searches of the *cox1* sequence in the non-redundant NCBI database to retrieve closely-related sequences. We aligned all the *cox1* sequences using the online version of MAFFT v. 7 (Katoh et al. 2019) available at https://mafft.cbrc.jp/alignment/server/. We estimated a maximum likelihood phylogeny from the resulting alignment with IQ-TREE v. 2.2.3 (Minh et al. 2020) and parameters -m MFP -seed 1234 -b100. Known karyotype numbers were obtained from the literature (Galvão et al. 2011; Prazdnikov 2023).

Synteny between the haplotypes was inferred by aligning the two scaffolds with minimap2 v. 2.28 (Li 2018) with parameters -X -N 50 -p 0.1 -c -B 4, and plotting them with the R package gggenomes v. 0.9.12.900 (Hackl et al. 2023). Only alignments larger than 5 Kb were considered for clarity. Alignments between TH1 and fGobFla1 were produced with the NUCmer program from the MUMmer package v. 3.23 (Kurtz et al. 2004) with parameters -b 200 -c 65. The scaffolds of the TH1 primary assembly were flipped to match the direction of fGobFla1 for downstream analyses.

#### 2.2 Identification of the Sex Determining (SD) region

##### ddRAD

For details regarding the ddRAD sequencing and initial Single Nucleotide Polymorphisms (SNPs) calling, we refer to Martinossi-Allibert et al 2025b, where the data was used to infer population structure of *P. flavescens* in Scandinavia. Briefly, we obtained ddRAD reads from 90 adult males and 90 adult females collected along the Norwegian coast in the summer of 2022 (Lat. 58N to 68N). Sex was assigned phenotypically in the field based on fin morphology and the presence of mature eggs in the case of females. In order to identify sex specific alleles, the ddRAD reads were aligned to a reference assembly, and variant calling was done with the Stacks pipeline (Catchen et al. 2013). When the project started there were no nuclear genomic resources available for *P. flavescens*. Hence we initially used the genome assembly of *P. minutus* as reference. The pipeline was re-run with the TH1 reference for other analyses unless otherwise stated. Association of variants with phenotypic sex was tested with PLINK 1.9 (Purcell et al. 2007, Chang et al. 2015), using the –assoc fisher option, and a group of 260 SNPs were identified. In addition, the ddRAD reads were processed with the RADsex pipeline (Feron et al. 2021) in order to identify sex-specific markers, using default settings and threshold depth of 1, supported by the *sgtr* package (Feron 2025) in R (R core team 2025).

##### Male-specific k-mer analyses

The cleaned forward and reverse Illumina reads were merged in a single file and fed to Jellyfish v. 2.2.10 (Marçais & Kingsford 2011) with parameters count -C -m 21 -s 100M to produce a k-mer count table per sample. After extracting a histogram table (parameter histo), we dumped the count table (parameter dump) with a minimum count of 2 per k-mer. We used the program Kpool v. 3.2.4 (Thompson et al. 2021) to merge (with kpool merge) the tables of same-sex individuals, producing a single count table per sex. We then merged the male and female tables into a single table (also with kpool merge) and estimated the male-specific k-mers with parameters -s M --min-het 30 --max-hom 5 based on the mean depth of coverage of all four samples. The k-mers with counts larger than 120 were removed as they could represent repeated (e.g., paralogs, transposons, etc.) sequences. The surviving k-mers were transformed into a fasta file and mapped to the two haplotype assemblies of TH1 using BWA with the commands aln and samse. We removed any k-mers that had secondary mappings or mistmatches and produced a table of coordinates for the unique-mapping k-mers. Any overlapping k-mers were merged to produce a BED file of male-specific genomic tracks. We used BEDtools v. 2.31.1 (Quinlan & Hall, 2010) to estimate the coverage of male k-mers in non-overlapping windows of 50 Kb.

To test whether specific scaffolds were enriched in male-associated k-mers, each window was classified as male-associated if its male k-mer coverage exceeded 0.05 (an arbitrary threshold chosen based on the coverage distribution). Enrichment was assessed using Fisher’s exact tests, where for each scaffold we compared the number of male-associated and non–male-associated windows inside the scaffold to those in all other scaffolds of the same haplotype. P-values were adjusted for multiple testing using the Benjamini–Hochberg false discovery rate (FDR) procedure. Scaffolds were considered significantly enriched when FDR-adjusted p < 0.05.

#### 2.3 Annotation of the SD scaffold

To produce a basic repeat library, we ran RepeatModeler v. 2.0.4 (Flynn et al. 2020) with the parameters -LTRStruct -quick and the following dependencies: LTR_retriever v. 2.9.0 (Ou & Jiang 2018), RMBlast v. 2.14.1+ (Smit & Hubley 2025), RepeatScout v.1.0.6 (Price et al. 2005), RECON v. 1.08 (Bao & Eddy 2002), MAFFT v. 7.526 (Katoh et al. 2019), GenomeTools v. 1.6.1 (Gremme et al. 2013), Tandem repeats finder (TRF) v. 4.10.0-rc.2 (Benson 1999), CD-HIT v. 4.8.1 (Fu et al. 2012), RepeatMasker v. 4.1.5 (Smit & Hubley 2025), and NINJA v.0.97-cluster_only (Wheeler 2009). To minimize computational resources, as input we only gave the largest scaffold in the primary assembly (scaffold ptg000015l, 38.8 Mb, corresponding to most of chromosome 4) and the two versions (hap1 and hap2) of the scaffold enriched for sex-specific markers.

To characterize the satellite DNA (satDNA) landscape, we concatenated the two phased assembly versions produced by Hifiasm (hap1 and hap2) into a single fasta file and ran TideCluster v. 1.6 (Novak 2025) with parameters -m 1000 -M 30000 --tidehunter_arguments ’-p 40 -P 5000 -c 5 -e 0.25’. We extracted the resulting superfamily monomers inferred by TideCluster, duplicated them, and created a library of satDNA families in fasta format using the script TideCluster2RM.py v. 1.1 available at https://github.com/SLAment/Genomics.

The RepeatModeler and TideCluster libraries were processed to remove potential protein-coding gene families using the Snakemake pipeline available at https://github.com/NBISweden/repeatlib_filtering_workflow. Briefly, transposon-related proteins were removed from the UniProt/SwissProt collection (reviewed proteins; downloaded on 2025-04-26) with TransposonPSI v.1.0.0 (https://transposonpsi.sourceforge.net), and this cleaned protein collection was used as query in BLAST searches against the RepeatModeler and TideCluster libraries. The families with hits were removed using ProtExcluder v. 1.2 (Campbell et al. 2014). The final TE and satDNA libraries were used to annotate the cleaned primary assembly, the hap1 version of the SD scaffold, and the reference fGobFla1 with RepeatMasker v. 4.1.8 (Smit & Hubley 2025).

We produced a basic protein-coding annotation with the online tool of Augustus v. 3.3.3 (Stanke et al. 2006) and the zebrafish training parameters, allowing for partial genes, and without additional hints or UTR prediction (the input assemblies were unmasked). We filtered out gene predictions that overlapped with any of the five most common satDNA families in the SD region (termed “TRC_1”, “TRC_3”, “sf10 TRC_75”, “sf11 TRC_42”, “TRC_105” in our library), as they tend to create low-complexity “proteins” if the repeat units happened to be in-frame. Remaining protein models were subjected to functional annotation using InterProScan v. 5.75-106.0 (Jones et al. 2014).

We manually examined any regions where the two females with Illumina data had zero coverage relatively to the Y haplotype (see Results), in search of male-specific sequences. Nearly all such regions were relatively small (less than 3 Kb) and contained repeats and protein-coding genes associated with transposable elements, such as transposases, RNase H, integrases, reverse transcriptases, and (retro) virus-related genes. However, we discovered a small gene model with three exons homologous to the C-term of the anti-Mulllerian hormone receptor type 2 (*amhr2*) gene. To improve on this gene model, we did blastn searches with the BLAST v. 2.15.0+ package (Altschul et al., 1997) on transcripts from testes and ovaries of *P. minutus* (Leder et al. 2021). We chose a variant with a coding sequence (CDS) that closely matched the *amhr2* homologs of the catfish family (Wen et al. 2022), corresponding to nine exons. We then used the *P. minutus*’ *amhr2* sequence to find and manually annotate homologs in *P. flavescens* while assuming canonical splicing sites (5’ GT - AG 3’). We found a single copy in the chromosome 10 of fGobFla1, and two in the TH1 assembly when including the haplotype 1 version of chromosome 16, both with eight exons. We failed to find one intermediate exon, corresponding to the end of exon 4 and the entire exon 5 of the catfish *amhr2* homolog (Wen et al. 2022). The missing sequence in *P. flavescens* is a fast-evolving area of the protein and might be found once transcriptomic data becomes available.

We confirmed that the copy of *amhr2* in chromosome 10 is ancestral by using it as query in BLAST searches within Genomicus v. 2021-08-15, database v. 110.01 (Nguyen et al. 2021). The search revealed two relatively conserved genes flanking *amhr2* in the ancestor of Actinopterygii (Ensembl codes ENSMAMG00000010416 and ENSMAMG00000010349 of the Zig-zag eel *Mastacembelus armatus*). We then performed tblastn searches using the *M. armatus* sequences on the TH1 assembly and found that their orthologs locate in chromosome 10, flanking *amhr2*.

#### 2.4 Diversity estimates

To estimate the heterozygosity of the individuals sequenced with Illumina technology, we called biallelic SNPs along chromosomes 15, 16, and 17, to the exclusion of other scaffolds to reduce computation time. Specifically, we mapped the cleaned Illumina reads using BWA and Picard as above using the fGobFla1 assembly as reference. We subsetted the BAM files by chromosome and used them as input for HaplotypeCaller of GATK v. 4.5.0.0 (McKenna et al. 2010). The resulting gvcf files were then used for joint variant calling with GenotypeGVCFs, applying the flag -all-sites. Sites (invariant and variable) with less than 20x or more than 300x were removed. Variant sites were further filtered out with the following conditions: QD < 2.0, FS > 10.0, ReadPosRankSum < -3.0, MQRankSum < -6.0, SOR > 3.0, and MQ < 40.0. Any site overlapping with annotated satDNA or other repeats were removed using BEDtools intersect. We used pixy v. 2.0.0.beta6 (Korunes and Samuk, 2021) to calculate heterozygosity (equivalent to π per individual) in non-overlapping 50 Kb-long windows. Windows with less than 10 Kb genotyped sites were treated as missing data.

#### 2.5 Genotype by sequencing

We used genotype by sequencing to determine the sex of individuals with no morphological signs of sex: eggs and larvae collected in artificial nests during the 2022 breeding season, and adults without visible secondary sexual characters, also captured during the 2022 season. To this end, we selected nine sex-linked loci from the Stacks results. Primers were designed using Primer3plus (Untergasser et al. 2012), and the Multiple Primer Analyzer from ThermoFisher was used to control for the possibility of cross-dimer formation (**Table S1**). DNA was extracted from 285 eggs, 300 larvae, and 96 adults for which phenotypic sex could not be established. The eggs and larvae were sampled uniformly from three of the six study populations (Kristineberg, Ringstad, and Hitra), from five nests or clutches in each population, with 29 to 66 individuals sampled in each nest; a clutch usually has mostly one father but often several mothers. The unsexed adults all originated from the northern population of Ringstad, since they predominantly occurred in that population. In addition, we extracted DNA for 10 phenotypic males and 10 females, to act as control. As for the ddRAD original markers, the choice of the sex-specific loci predates the production of the *P. flavescens* genome assemblies and hence relied on the *P. minutus* reference. The mapped reads were used for variant calling with the Stacks pipeline as described above.

To evaluate the marker quality, we calculated the major allele frequency (MAF) in the read counts of each marker and examined their distribution (**Figure S1**). By design, the nine chosen markers should be homozygous in the females and hence should have a MAF close to 1. Males are expected to be heterozygous and thus should have a MAF centered around 0.5. One marker was discarded based on having mostly missing data and three markers were discarded for having frequency modes of intermediate values (∼ 0.85) in the males, which might indicate hidden paralogy (**Figure S1a**). Based on the MAF distributions of the controls and other samples, a given genotyped individual (eggs, larvae, or adults of unknown sex) was considered genetically female if the average MAF of all five remaining markers was >0.9, male if <0.65, and otherwise was left as undetermined (**Figure S1b**).

The phenotyping of controls had very high certainty, since all 10 females were carrying mature eggs, and all 10 males exhibited typical male fin morphology and were collected guarding nests (male-specific activity). However, one of these 20 control individuals, a phenotypic male, was assigned the female genotypic sex with high confidence (**Figure S1**), potentially due to mislabeling or recombination between the marker and the SD locus.

## 3. Statistical analyses

### 3.1 Analysis of sex ratio data

#### 2022 observation transect data

We analysed the temporal trajectory of ASR during the 2022 breeding season with a generalised mixed-effects model using the *lme4* package (Bates et al. 2015) in R (R Core Team 2025). In this binomial model, the response variable is the count of males and females of each observation transect. The fixed effects are the date (day of the year scaled), the date squared, and the population of origin, as well as the interactions of (i) population of origin and date and (ii) population and date squared. We allow random slopes for each sub-location by the addition of a random effect.

#### Sex ratio of eggs, larvae and unsexed adults

We analyzed the effect of latitude on sex ratio of eggs and larvae in the same binomial generalised model, with the count of genetic males and females as the response variable, and latitude and type of sample (egg or larvae) as fixed effects. The interaction between latitude and sample type, not being statistically significant, was not included in the final model.

The sex ratio of unsexed adults in the population of Ringstad was assessed with a generalised binomial model, with the count of genetic males and females as the response variable, and time of the season (early, mid, late) as the only fixed effect.

## Results

### 1. Genetic sex determination in *P. flavescens*

#### 1.1. First genome assembly of a male two-spotted goby

At the start of the project, no genomic resources of *P. flavescens* were available to investigate the genetic basis of SD. Thus, we produced a genome assembly of a male individual from Trondheim, Norway, called TH1, based on PacBio HiFi reads. After cleaning, the primary TH1 assembly contained 411 nuclear scaffolds, with a total genome size of 871.45 Gbp, an N50 of 22.88 Mbp, and an average depth of coverage of 236x. Around the same time, a more continuous *P. flavescens* assembly was published as part of the ATLASea programme (https://www.atlasea.fr). The sequenced individual came from France (code fGobFla1) and is of unknown sex but contains the 23 chromosomes expected from cytogenetic work (Klinkhardt 1992). The fGobFla1 nuclear assembly is 826.72 Gbp and highly collinear with our assembly, with the exception of two translocations involving chromosomes 3, 7, 11, and 20 (**Figure S2**). These structural variants might be misassemblies or true structural differences. Hence, we considered the French individual as the reference genome of the species, but used the TH1 male assembly whenever haplotype differences were relevant.

A maximum likelihood phylogeny of *cox1*, a common DNA barcode for animals, confirmed that all these sequenced *P. flavescens* individuals form a monophyletic clade of nearly identical sequences (**Figure S3**). Surprisingly, the sequences labeled as *P. minutus* were polyphyletic, which might explain the high karyotypic diversity previously reported for this species (Klinkhardt 1992).

#### 1.2. The two-spotted goby has an XX/XY determination system

We tested for association of allele frequency with sex using the SNPs data obtained from the ddRAD sequencing of 178 adult individuals collected across the latitudinal range of our study. This analysis revealed a group of SNPs, present in both sexes, but with intermediate frequency in males and close to fixation in females (**Figure S4**). To obtain a more complete set of sex-linked markers, we used the dedicated RADsex pipeline with mapping to the TH1 assemblies, which showed a group of markers found disproportionately in males (**Figure 2a**). The combination of male-specific markers and intermediate allele frequency of sex-linked SNPs in males is consistent with an XX/XY sex determination system.

**Figure 2.**
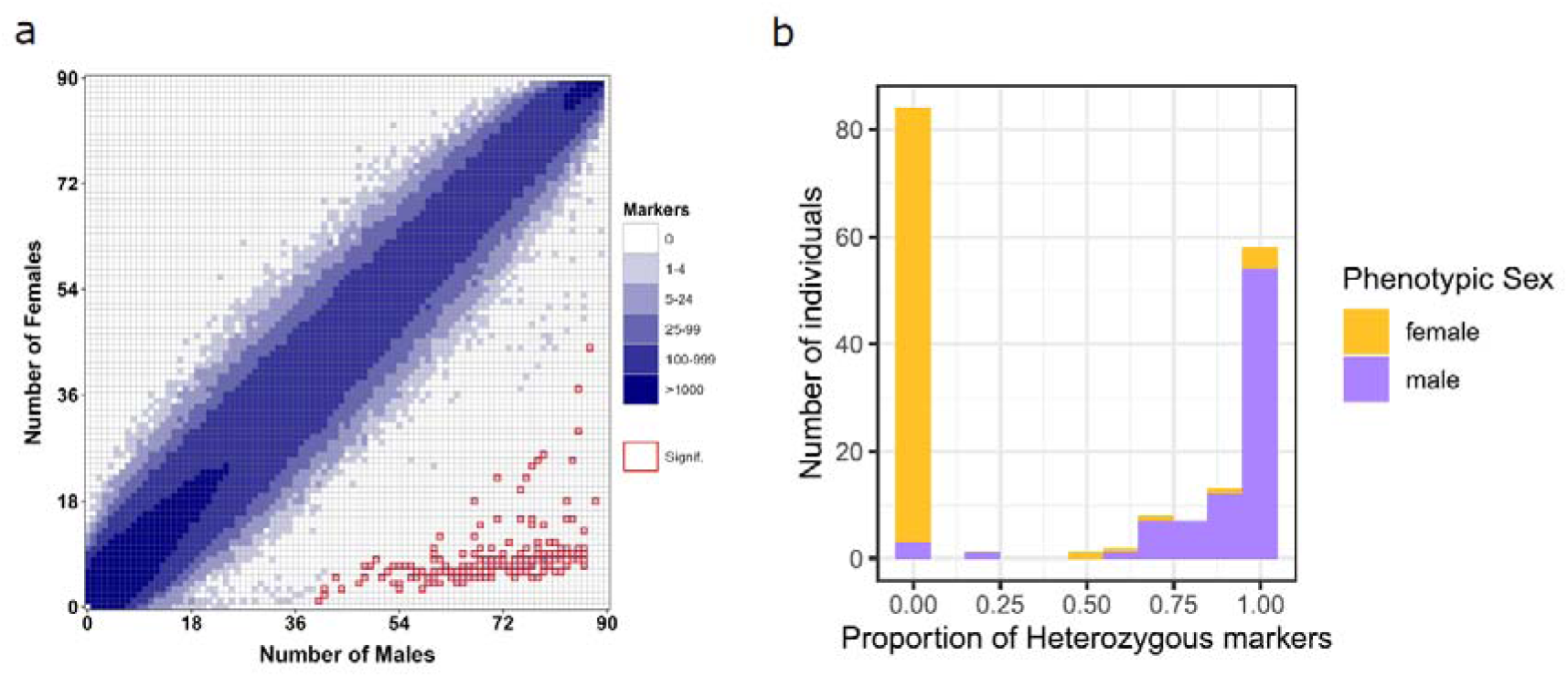
Sex-specific occurrence of all markers from the ddRAD dataset in the 89 females and 89 males *P. flavescens* (a), and proportion of the nine candidate markers for sex identification that are heterozygous among the same individuals (b), indicating a XX/XY sex determination system. *In (a) the position of a cell indicates the number of males and females in which a marker was found, and the colour of a cell indicates the number of markers corresponding to that combination (no depth filtering). The red outlines indicate significant association with sex. The pipeline identified 203 significantly male biased markers. In (b), the number of individuals for different proportions of heterozygous markers among the nine markers selected for sex identification. The colour of the bars represents the phenotypic sex. The expectation with an XX/XY sex determination system would be that all females are homozygous and all males heterozygous, if the markers are within a region perfectly linked to sex*.

We note that the association of phenotypic and genetic sex was not perfect among the 178 ddRAD sequenced adult individuals. We selected nine markers among the SNPs identified from the ddRAD data, to be used for identification of genetic sex (used in Results 2.1). Most phenotypic females (81) were homozygous for these polymorphisms, as expected for a XX/XY system. However, eight females had half or more of their markers heterozygous, including four phenotypic females with all markers heterozygous (five to eight out of nine successfully sequenced **Figure 2b**). Conversely, 81 males had more than half of their markers heterozygous, including 54 with all markers heterozygous. Four phenotypic males had more than half of their markers homozygous, including three having all markers homozygous (five to seven out of nine successfully sequenced, **Figure 2b**). Thus, the association of phenotypic and genetic sex is found in about 95% of cases, with exceptions in both sexes.

#### 1.3. The XX/XY system is defined by a small chromosomal region with little differentiation

We found that all RADsex markers associated with phenotypic sex map to a single specific scaffold around 5 Mb long in each of the haplotype-resolved TH1 assemblies. These two haplotypes are homologous and correspond to the distalend of chromosome 16 in the French reference fGobFla1 (**Figure 3** and **Figure S5**), as confirmed by the presence of telomeric repeats on their right edge. While highly collinear, the two TH1 haplotype versions of this region are differentiated by large tracks of various satellite DNA (satDNA) families (**Figure 3**). The observed satDNA families are not specific to this particular scaffold, as satDNA blocks were found at the end of chromosome arms across the *P. flavescens* genome (**Figure S6**).

**Figure 3.**
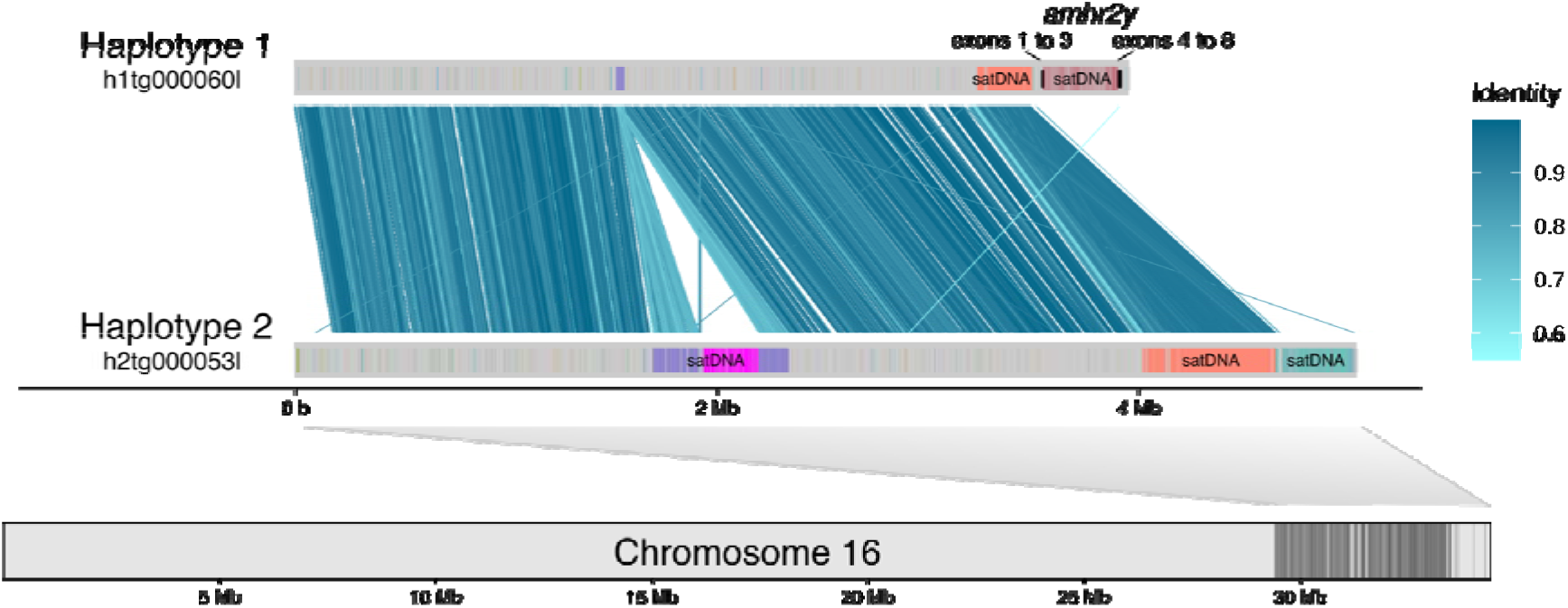
Scaffold alignment between the alternative haplotypes of TH1 matching the right end of chromosome 16. *The scaffolds are represented as horizontal gray bars, with satDNA families marked within the scaffolds using different colors. Alignments smaller than 5 Kbp were ignored for clarity. The dark gray shading in chromosome 16 corresponds to direct alignments between the haplotype 2 of TH1 and the reference genome fGobFla1. The location of the* amhr2y *exons within haplotype 1 is marked with vertical black lines*.

In an effort to determine which haplotype corresponds to either the X or the Y chromosome, we sequenced two phenotypic males and two phenotypic females with Illumina technology. The individuals were chosen to represent the extremes of their geographical distribution, with one sex pair belonging to the southern location of Kristineberg and the other to the northern location of Helligvær (**Figure 1a**). From the Illumina reads we calculated the coverage (fraction of bp) of male-associated k-mers in windows of 50 Kb along the haplotype resolved TH1 genome. The small number of samples unavoidably creates background noise, where any shared heterozygosity between the two males that is absent in the two females is identified as a male-specific K-mer. Despite these limitations due to low sample numbers, this strategy revealed a significant enrichment (FDR-adjusted *p* = 5.17e-25) of male-associated k-mers exclusively in the haplotype 1 version of the right end of chromosome 16; a similar pattern was not found on other scaffolds (**Figure S7**). By contrast, the haplotype 2 version had a noticeable absence of male-associated k-mers (**Figure 4**). This finding suggests that the haplotype 1 of TH1 can be interpreted as the Y chromosome and haplotype 2 as the X chromosome (i.e., the gametologue of the Y). Moreover, while the assembly of haplotypes is not perfect with PacBio data alone (Cheng et al. 2021), the spatial clustering of windows with relatively high male-associated kmer coverage suggests that the Y haplotype was correctly reconstructed, at least within the area flanked by satDNA blocks. Herein, we define the SD region as the 2.4 Mb chromosomal area covered by RADsex markers and male-associated kmers on haplotype 1 (**Figure 4**).

**Figure 4.**
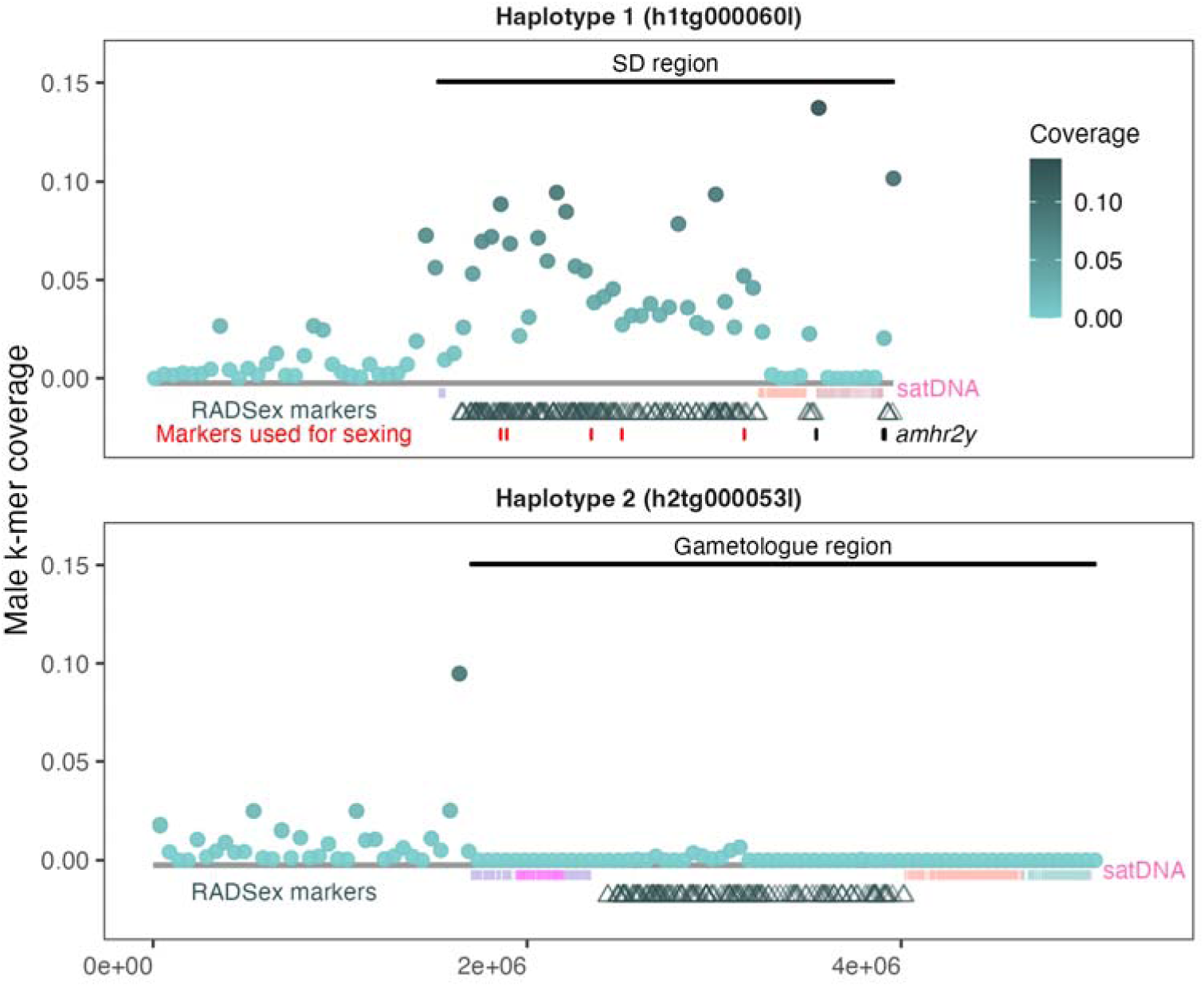
Distribution of male k-mers and RADsex markers in the two sex-linked scaffolds from the haplotype-resolved TH1 assemblies, confirming haplotype 1 as the Y-specific sequence. *Coverage of male k-mers (fraction of bp) was calculated in non-overlapping 50 Kb-long windows. The annotation of satDNA is highlighted below each scaffold (only five most common families shown for clarity). Notice the scaffolds themselves are not aligned to each other. The markers used for sexing eggs and larvae are highlighted in red for the haplotype 1. Likewise, the exons of the* amhr2y *gene are marked in black*.

As a proxy of divergence between haplotypes, we further calculated heterozygosity along the genomes of the four Illumina samples using fGobFla1 as reference. As expected, the two females are significantly less heterozygous in the sex-linked region than the males (**Figure 5**). However, this region is not particularly heterozygous in the males either, compared to other chromosomal areas. Rather, there is considerable variation in the distribution of heterozygosity along the chromosomes of *P. flavescens* (**Figure 5a**), which might reflect variation in the recombination landscape. Thus, despite the clear association of sex with the SD region, there seems to be no obvious differentiation between the X and Y haplotypes relative to the rest of the genome. As a note, the male of Helligvær (PflaHELEm) is considerably more heterozygous in other parts of the genome than the other individuals, including the female of its own population (**Figure 5c**).

**Figure 5.**
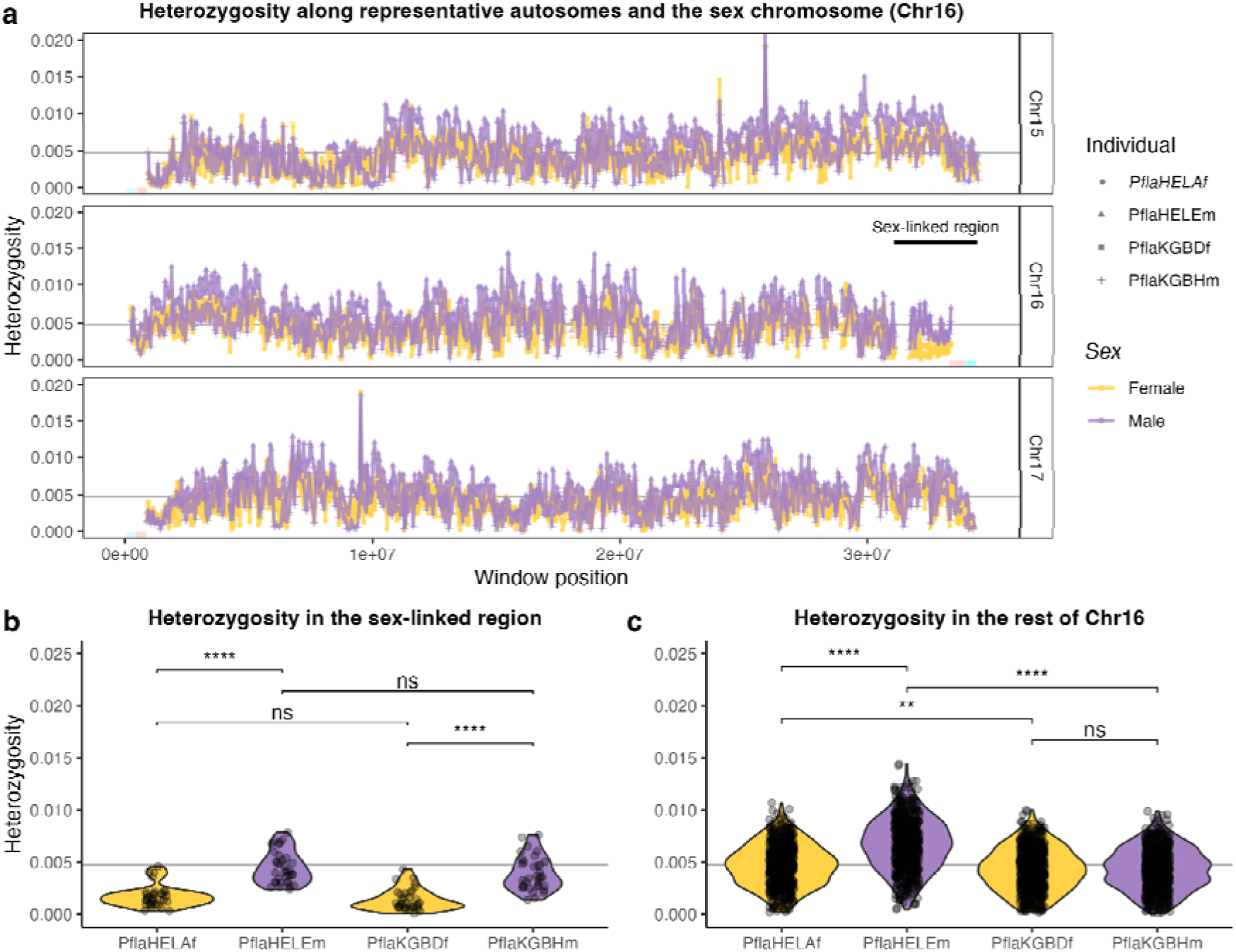
Genetic diversity in the sex chromosome and selected autosomes of *P. flavescens* individuals sequenced with Illumina technology. *(**a**) Heterozygosity calculated in non-overlapping 50 Kb-long windows along the chromosome containing the sex-linked region (chromosome 16) and two other chromosomes of similar size. (**b**) Violin plots depicting the distribution of window heterozygosity in the sex-linked region and (**c**) in the rest of chromosome 16. The five most common satDNA families are highlighted with color ticks on the x-axis of each chromosome. The average genome heterozygosity of the four individuals (*π *= 0.00478) is marked with a gray horizontal line. Significance determined by Wilcoxon rank sum tests with Bonferroni correction (****: p <= 0.0001; **: p <= 0.01; ns: not significantly different)*.

#### 1.4. The subtelomeric region of the SD region contains a translocated paralog of *amhr2*

In agreement with their high collinearity and similarity, we found no conspicuous coverage differences between the two sex-linked haplotypes apart from the satDNA areas (**Figure S8** and **S9**). Yet, one exception represents a strong candidate for the master sex-determining (MSD) gene: a copy of the anti-Mulllerian hormone receptor type 2 (*amhr2*) gene in the subtelomeric end of the Y haplotype (**Figure 3**). The *amhr2* gene is part of the TGF-β signaling pathway, whose members have evolved MSD roles multiple times across teleosts (Pan et al. 2021). The *amhr2* gene in particular acts as the MSD in two unrelated fish families, Tetraodontidae (pufferfishes) and Pangasiidae (catfishes) (Kamiya et al. 2012; Wen et al. 2022).

In the catfish family in particular, the ortholog of *amhr2* underwent a duplication, producing a N-terminus truncated version that became the MSD gene, while the original copy remained intact (Wen et al. 2022). Similarly, we found the ancestral copy of *amhr2* in chromosome 10 in *P. flavescens*. This *amhr2* gene neatly aligns to that of the Pangasiidae, including the N-term transmembrane domain and the C-term kinase domain (**Figure S10**). The *amhr2* duplication in the Y haplotype, herein *amhr2y*, retains the same exons as its paralog in chromosome 10, with little differentiation (90.7% amino acid identity). However we failed to find the starting codon without further evidence from transcriptomic data, suggesting 1) the gene model is incomplete, 2) its a pseudogene, or 3) it evolved a new starting codon up or downstream. Remarkably, the first exons of *amhr2y* are separated by nearly 350 Kb entirely composed of a single satDNA family (**Figure 3** and **4**), defined by GGTCTCTCTCT units and close variations. Given the high amino-acid identity and conserved exonic structure, we interpret these findings as a recent duplication of *amhr2* into the Y haplotype, followed by a dramatic intronic satDNA expansion. Recent translocation of this gene into this genomic region may explain why extended differentiation between the putative X and Y haplotypes is not observed. It remains to be confirmed if *amhr2y* has acquired a secondary start codon downstream, emulating the loss of the N-term in the catfish family homolog, and if it truly functions as the two-spotted goby’s MSD that triggers testicular development in males.

In summary, our findings point to an XY sex determination system with very little sex-chromosome differentiation. The identification of a putative SD locus at the very end of chromosome 16, and the strong association between phenotypic sex and genotype, enables us to estimate the sex ratio in development stages for which phenotyping is not possible.

### 2. Sex ratios in *P. flavescens* populations

#### 2.1. No overall bias in the sex ratio of eggs and immature adults

Having selected nine SNPs strongly associated with phenotypic sex based on the ddRAD data (see Results 1.2), we assessed the genetic sex of 585 eggs and hatched larvae collected in artificial nests placed in natural conditions across the latitudinal range of the study, with the aim of detecting departures from the 1:1 sex ratio and eventual latitudinal effects on sex ratio close to the hatching stage. Based on the success of their amplification and sex discrimination power, we restricted our analysis to five of the nine markers (**Figure 4** and **Figure S1**). Considering these markers, the genetic sex ratio of eggs and larvae across all study populations, expressed as the proportion of males, was: SR = 0.47, 95%CI [0.43;0.51], and for each of the three populations from South to North: Kristineberg = 0.43 , 95%CI [0.36;0.50] , Hitra = 0.46, 95%CI [0.38;0.53], Ringstad = 0.52, 95%CI [0.45;0.59] (**Figure 6a**). A positive effect of latitude on the proportion of males among eggs and larvae was close to the significance threshold (*p* = 0.07, **Table 1**).

**Figure 6.**
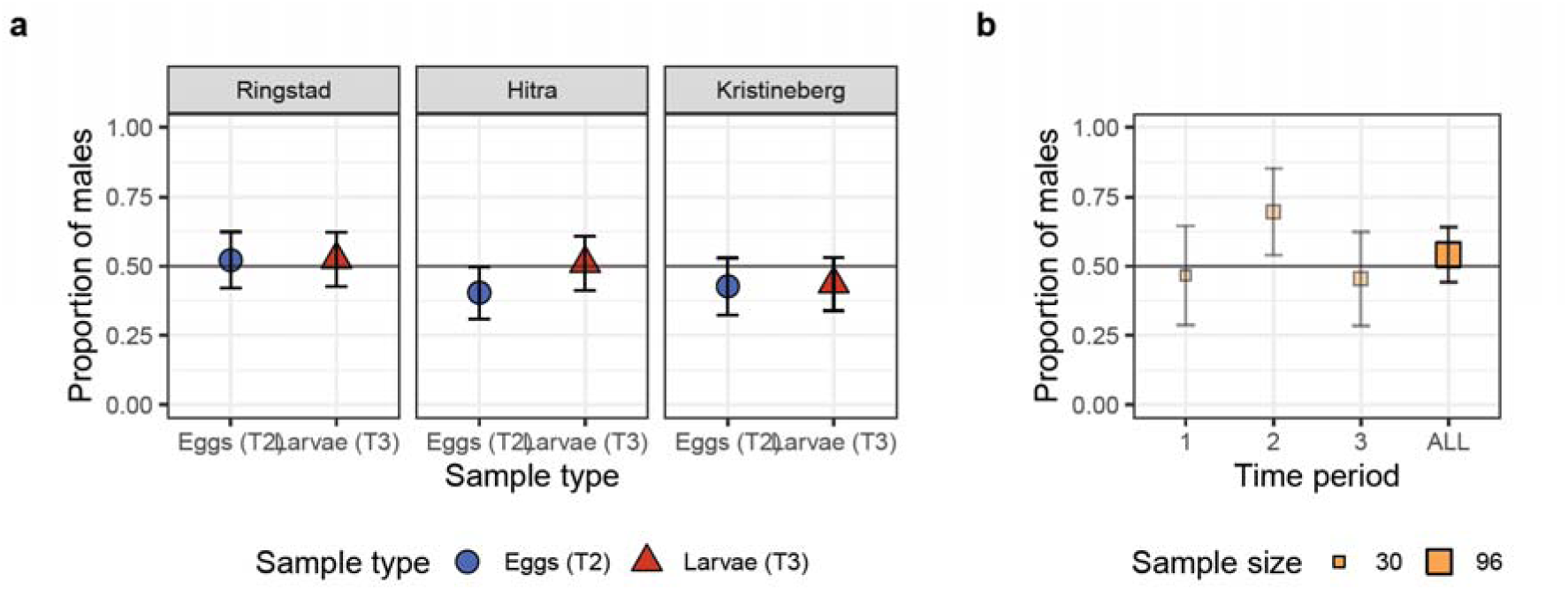
Genetic sex ratio of *P. flavescens* (a) eggs and larvae from three populations and (b) unsexed adults from one population of the 2022 field season. *The proportion of males identified by genotyping is given for each sample. Error bars represent a 95% confidence interval calculated according to the Normal approximation method. In (a), the range of sample sizes for each shape is: 89-104. Eggs were collected in peak breeding season (T2, June 2022) and the just hatched larvae in late season (T3, July 2022). In (b), ninety six immature 1-year-old individuals from the Ringstad population were genotyped. The range of sample sizes for each time period is: 30-33*.

**Table 1.** Type II anova table for the binomial generalised linear model fitting the effect of Latitude and sample type on the sex ratio of eggs and larvae.

| Effect | Chi square | df | p |
| --- | --- | --- | --- |
| Latitude | 3.26 | 1 | 0.073 |
| Type (egg/larvae) | 1.03 | 1 | 0.311 |

In addition, when examining the sex ratio at the level of each clutch or nest, we could not exclude that some clutches (2 out of 15) may have had female biased sex ratios (**Figure S11**).

Individuals from northern populations frequently postpone their reproduction to the second year of life, to compensate for limiting growth conditions (Martinossi-Allibert et al. 2025b). The sex ratio in this class of individuals could inform on eventual sex-specific mortality during the first year of growth. In 2022, we collected sexually immature 1-year-old individuals (age determined by size class) in the northernmost population of Ringstad, where they represented 22% of the adult population (Martinossi-Allibert et al. 2025b). Using the same markers as for the eggs and larvae, we assessed their genetic sex. The overall sex ratio of immature 1-year-old fish, expressed as the proportion of males, was: SR=0.54, 95%CI[0.47,0.61] (**Figure 6b**). Moreover, the sex-ratio of immature 1-year-olds did not differ across sampling times (X^2^=5.0, df=2, *p*=0.082, **Figure 6b**). These results indicate that males and females likely postpone reproduction in similar proportion in the northernmost population, and do not appear to suffer sex-biased mortality.

#### 2.2. The Adult Sex Ratio is generally female-biased across latitude and time

ASR in 2022 was generally female-biased, with the exception of the early season sampling in the two northernmost populations (**Figure 7** and **S12**). ASR generally decreased over time across all populations (**Table 2** and **Figure 7**). However, populations differed significantly in their trajectories, and some do not show a constant decrease over time (see for example the Arendal population in **Figure 7** and **Figure S11**).

**Figure 7.**
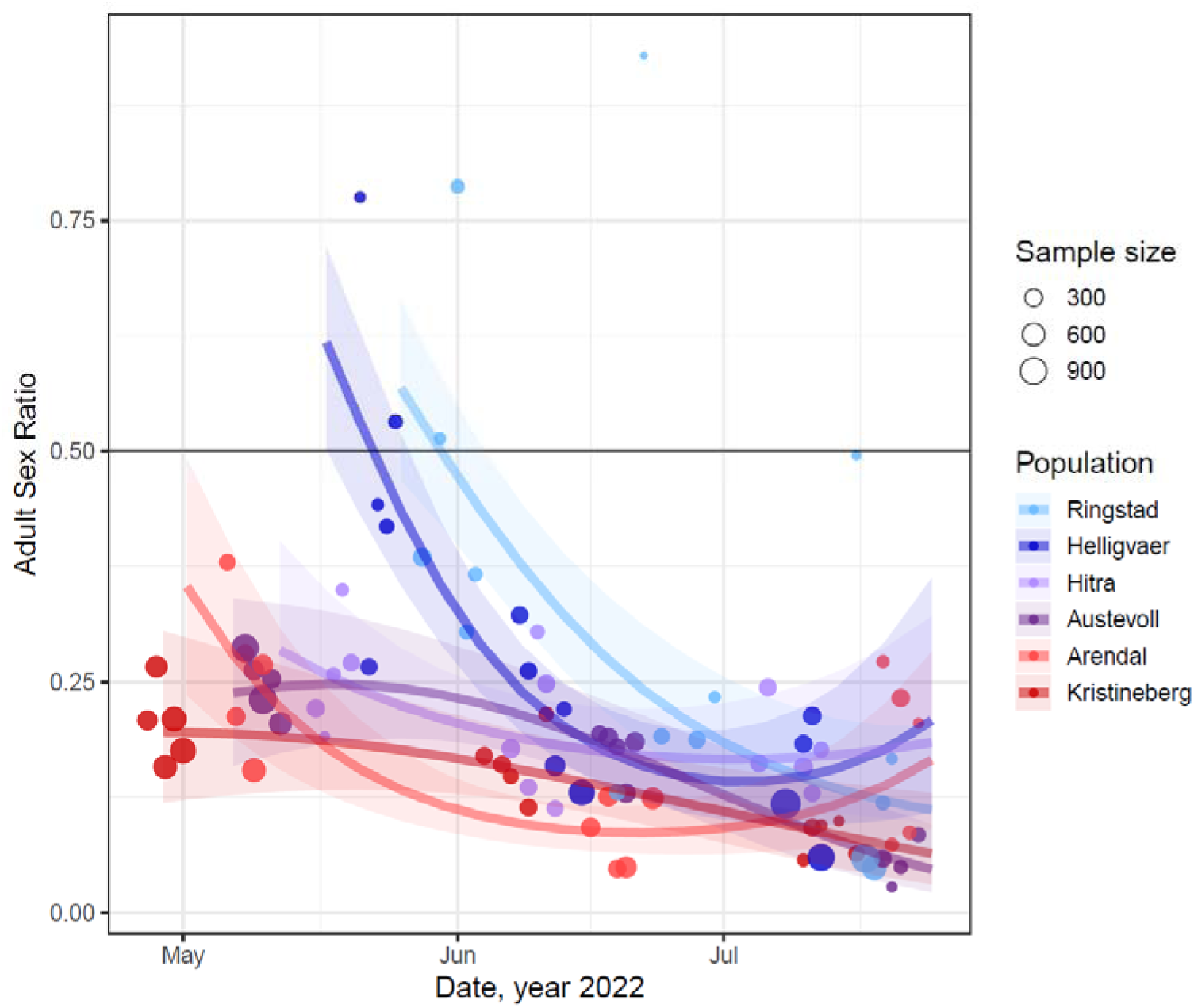
Adult sex ratio (ASR) from observation transects in the six Scandinavian populations during the 2022 breeding season. *Individuals were counted and sexed by snorkeling observers along transects. Each population is represented by five transects, and each transect was sampled three times (early, mid- and late- season). ASR is calculated as the proportion of males: m/(m+f). Each dot represents a sampling occurrence for a transect, with the size of the dot proportional to the total adult census. The curves for each population represent the fit from a binomial model (see Table 1), with 95% confidence intervals encompassed by the shaded area around each curve. The horizontal black line indicates a balanced sex ratio*.

**Table 2.** Type III ANOVA for the binomial mixed-effects model of the temporal trajectories of ASR in the 2022 observation transect data.

| Effect | Chi square | df | p |
| --- | --- | --- | --- |
| Intercept | 143 | 1 | <0.001 |
| Date | 4.77 | 1 | 0.029 |
| Date^2 | 3.74 | 1 | 0.053 |
| Population | 50.8 | 5 | <0.001 |
| Population:Date | 21.8 | 5 | <0.001 |
| Population:Date^2 | 112 | 5 | <0.001 |

#### 2.3. Effect of collection method on ASR

Because ASR can be sensitive to the sampling method, we compare the 2022 snorkeling observation data (observer phenotyping fish freely swimming in the water) to catch data (observer phenotyping fish in hand on land) from the same sampling effort. Both samplings were performed on the same day, in the same environment; catch data should be more robust to phenotyping errors, but could be more sensitive to selection bias, especially because several experimentators were involved in the capture of the fish. Nevertheless, the two sampling methods show similar trends and are well correlated at the site level (correlation coefficient=0.71, 95% CI [0.59,0.80], **Figure S13**).

To give more perspective to the 2022 ASR field data, we obtained unpublished data from previous *P. flavescens* sampling efforts, in 2007, 2008, 2010 and 2019 in three locations in Scandinavia, two of which are close to the populations in the 2022 sampling (Kristineberg and Austevoll/Bergen). The third population is located on the Finnish side of the Baltic Sea (**Figure 1**). All these samplings targeted the breeding habitat. Some of these ASR were estimated by snorkeling observation transect, similar to the 2022 sampling, but others were beach seine samplings (fishing net operated from shore with the aid of a small boat). As in 2022, we observe a general female-bias but with some clear exception of balanced or even male-biased sex ratio, not always at the beginning of the breeding season (**Figure 8**).

**Figure 8.**
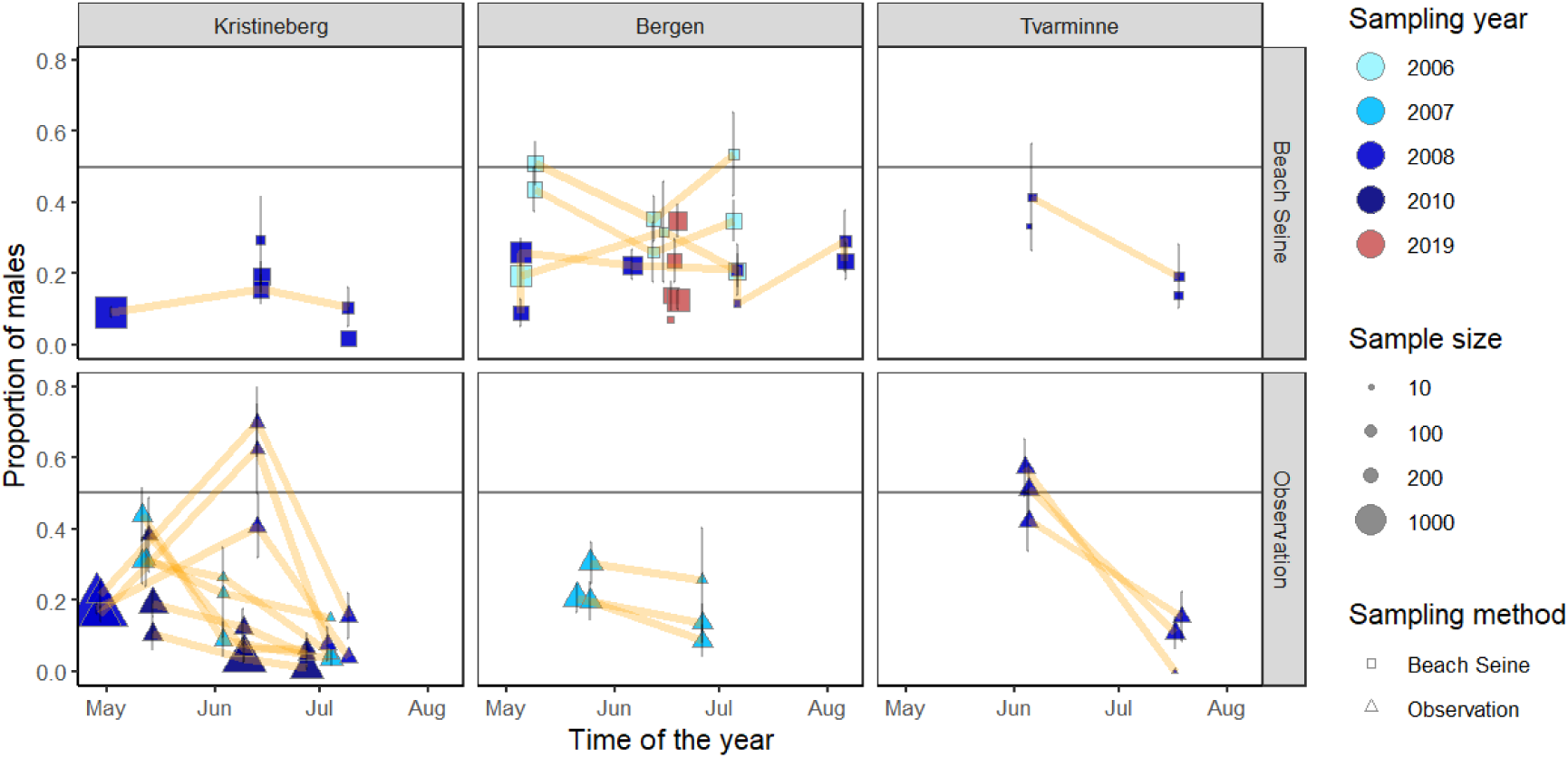
Adult Sex Ratio of *P. flavescens* in three different natural populations, over several sampling years and with different sampling methods. *Each panel is a sampling population for a given sampling method. Error bars represent 95% confidence intervals calculated according to the Normal approximation method. Data points without error bars did not fulfil the assumption for Normal approximation. The yellow lines connect samplings that were made in the exact same location within each population. Sampling method is indicated by the shape, sampling year by the colour, and sample size by the size of the dots*.

## Discussion

Our extensive sampling effort in 2022 revealed a consistently female-biased Adult Sex Ratio (ASR) in *P. flavescens* in the breeding habitat across the entire Scandinavian range. This observation is concordant with estimates from the southern part of the range from earlier publications (Forsgren et al. 2004), and we consolidated it with additional datasets from the period 2008-2019. At the same time, our genomic investigation demonstrated that sex determination is largely, if not completely, genetic in *P. flavescens*, and we could not detect any substantial departure from a balanced genetic sex ratio in early life stages. The explanation for a marked and geographically widespread female bias in adults is thus unlikely to be found in the sex determination system alone. Nevertheless, we want to highlight twoimportant points regarding these novel sex determination results: the possibility for environmental contribution to sex determination, and the robustness and limitations of our estimates of genetic sex ratio. Addressing these points is essential for interpreting the observed ASR trends and to place our findings within a broader ecological and evolutionary context.

The strong association between genetic markers and phenotypic sex, as well as the presence of a male-specific genomic region containing a gene whose homologs controls sex determination in at least two other fish groups (Kamiya et al. 2012; Wen et al. 2022), gives us high confidence that *P. flavescens* has an XX/XY sex determination system. Still, we cannot completely exclude the possibility of environmental contribution to sex determination. Partial ESD is common in fish: masculinization at high temperature is a frequent outcome, even in species previously identified as of purely GSD (Baroiller and D’Cotta 2016, Geffroy and Wedekind 2020, Lema et al. 2024). Mechanisms of partial ESD can be hormonal or epigenetic (e.g., Geffroy and Wedekind 2020) and do not seem to be associated with a particular genetic architecture of sex determination, be it chromosomal or polygenic (Valdivieso et al. 2022). In our ddRAD sample of 178 adults, the association between phenotypic and genetic sex was not complete, with about 5% mismatches in both directions. Such mismatches could stem from methodological limitations and are thus not conclusive evidence for ESD: first, phenotyping errors are possible in individuals with undeveloped secondary sexual characters, second the genetic markers of sex were obtained from reduced representation sequencing, and their linkage to the sex determination locus may not be perfect. Nevertheless, we acknowledge the possibility for environmental effects on sex determination, which would not be detected in the genetic sex of early life stages, but could still contribute to biases in the phenotypic ASR. In this case however, we have no indication that ESD could lead to a general adult sex-bias since mismatches occurred in equal numbers in both directions.

Another result that suggests the possibility of ESD is the ASR latitudinal pattern in 2022, with an absence of female bias in the two northernmost populations at the onset of the breeding season. This remains very indirect and inconclusive evidence for ESD, but it opens the possibility for a climatic effect on sex determination.

Because of the global sample size, we are confident that estimates of sex ratio in early life stages exclude genetic sex determination as the main driver of the strong female bias among adults. Statistical power to detect global effects large enough to explain the global ASR bias was high: 87% power to detect 0.1 deviation from a 0.5 sex ratio. Nevertheless, effects of smaller magnitude (below 0.1), or more localised (single population or single nest, implying reduced sample size), may remain undetected. In particular, there are two observations that point to weak effects of interest. First, the egg sex ratio had a female bias in two of the 15 clutches assessed, which could be the result of, for example, an X-linked meiotic driver introducing segregation distortion during meiosis or fertilization rate difference between the X and Y bearing sperm (e.g., Jaenike 2001). Second, the proportion of males among eggs and larvae could be increasing with latitude. These trends go in the direction of sex ratio patterns observed among adults, and thus are worth investigating further; even though such effects would remain of too small a magnitude to completely explain ASR patterns, they could contribute to them.

In 2022, as well as in the earlier data from the period 2004-2019, the proportion of males tended to decrease throughout the breeding season (May to July), with a few exceptions. This global trend, comparable to what was observed in Fries goby *Lesueurigobius friesii* in Scottish waters (Gibson and Ezzi 1978), is consistent with a higher mortality of males associated with the cost of nest brooding activities. In *P. flavescens*, the cost of nest guarding is expressed in reduced foraging opportunities and decreased body condition (Skolbekken and Utne-Palm 2001). The parasitic load of *P. flavescens* males in southern populations is generally high at the end of the breeding season (Utne-Palm, personal observations), a further indication of high male energy expenditure during that period.

While breeding related male mortality could explain the decrease of male presence over the span of the breeding season, we do not believe that it can account for female-biased ASR at the onset of breeding in early May. Since a vast majority of breeders in southern populations are 1-year-old fish at their first breeding attempt (Martinossi-Allibert et al. 2025b), breeding related mortality cannot have had an effect on ASR already. Moreover, a high impact of breeding-related mortality should result in a more pronounced female-bias among second-year breeders, but this is not what we observe. In the two northernmost populations (Helligvær and Ringstad), a large proportion of breeders are 2-year-old, and some likely breed two consecutive years (Martinossi-Allibert et al. 2025b): still, the ASR at the onset of breeding was less female biased in these populations than in southern ones in 2022. Thus, we are left with ASR patterns that do not match what could be expected from the sex determination system and mortality prior to the breeding period. Next, we consider potential limitations of ASR estimates.

It is generally challenging to measure ASR in wild populations because the sexes often adopt different lifestyles, which can lead to sampling biases (Ancona et al. 2017, Kappeler 2023). This is especially a concern in marine fish, where samples are dependent on area, season, and method of catch (Sadovy and Colin 1995, Kendall and Quinn 2013). ASRs reported for marine species vary widely (e.g. Clarke 1983, Oliveira et al. 2012), and only a handful of comprehensive studies are able to distinguish methodological artifacts from true demographic trends. For instance, when populations of the cobaltcap silverside *Hypoatherina tsurugae* in the Tokyo bay switched from balanced to male-biased sex ratio over a 3-year period, Miyoshi et al. (2020) showed that it was a demographic trend explained by effects of water temperature on the sex determination system. In the rainbow trout, a facultatively anadromous fish, males can be largely over-represented in the non-anadromous strategy, providing a striking example of how sex-specific life histories can result in locally biased sex ratios (Rundio et al., 2012). Studies in the black goby, *Gobius niger*, show a combination of sex-specific life history and sampling biases affecting sex ratio estimates. Aydın and Karadurmuş (2022) reported almost balanced sex ratios outside of the breeding season, but 80% males on average during the breeding season, close to shore where males establish nesting sites. At the same time using bottom trawling instead of gill nets, presumably less selective, Kasapoğlu (2016) reported female-biased sex ratios instead.

In the case of *P. flavescens*, we can narrow down possibilities thanks to the wealth of ASR data collected in the field using various methods. In 2022, a direct comparison of (i) population census in the water, and (ii) population census by individual catch and phenotyping on land, suggested that there is little scope for systematic phenotyping errors during observations or sex-bias during the catch. Putting in perspective the 2022 data with samples from the period 2004-2019, which include method (i) and (iii) population census by beach seine, further shows that the global female-bias of ASR is robust to sampling method.

Beyond this global female bias, however, all three methods gave sporadic observations of balanced or even male-biased ASR, in the same year and locations that were previously sampled as female biased. In our view, the fact that study sites can sometimes gain males over time is evidence supporting the migration of individuals, either across areas of the breeding habitat, or from outside the breeding habitat. Such plasticity in behavior would not be surprising since *P. flavescens* is a model system for the study of effects of the social environment (Amundsen 2018), and several studies have shown that courtship behavior can vary in response to the local sex ratio in the breeding habitat (Forsgren et al.2004, deJong et al. 2012, Martinossi-Allibert et al. 2025a).

If *P. flavescens* individuals have alternative habitats that they favor outside of the breeding period, as suggested by their disappearance from the shoreline in winter (Amundsen 2018), this offers a potential explanation for the general female-biased ASR as well as ASR fluctuations with latitude and over time. With the choice of two alternative habitats, individuals may base their decision to migrate on the prospects of successful breeding, weighed against the cost of staying in the breeding habitat (e.g., higher risk of predation and infection, higher food competition). If these costs and benefits are sex-specific, a sex-bias in habitat distribution would arise, and if these parameters vary over latitude and time, the ASR in the breeding habitat could fluctuate accordingly.

For each sex, the prospect of successful breeding is likely to be inversely proportional to the abundance of same-sex competitors in the breeding habitat. In *P. flavescens,* one male can provide care for the clutches of several females at once, meaning that prospect of breeding likely decreases faster in males than in females with the number of competitors. This asymmetry in the reproductive strategy, originating in the mode of parental care, could be enough to cause sex-specific habitat distribution. A latitudinal effect could arise due to the much shorter breeding season in Northern populations: a short time window eliminates the possibility for delayed breeding, so individuals in northern populations might attempt breeding even if competition is high. In southern populations, the long breeding season may instead allow for early and late breeder strategies with reduced competition.

The hypothesis of breeding related sex-specific habitat distribution is supported by our observation of a balanced genetic sex ratio in pre-breeding adult shoals in the northernmost population. If the female biased ASR is dictated by mating related decisions, it wouldn’t be apparent in non-breeding individuals, even if occasionally present in the breeding habitat. More latitudinal and temporal data of ASR in the breeding habitat, together with a better overview of *P. flavescens*’ range outside of the breeding habitat would be required to test this hypothesis further.

Beyond the drivers of ASR, our exploration of the genomic architecture of sex determination in *P. flavescens* yields evolutionary and mechanistic insights on the evolution of sex chromosomes. Early karyological work found no evidence of heteromorphic sex chromosomes in *P. flavescens* or in related gobies (Klinkhardt, 1992). Accordingly, we found that the SD region is relatively small (2.4 Mb) and displays low levels of divergence relative to its gametologue. It is unclear if closely related species share the same SD locus, as current genomic resources are fragmented and limited. On the other hand, our candidate MSD gene, *amhr2y*, represents a potential additional case of convergent evolution of SD architecture in fish. Teleosts ancestrally carry a single copy of *amhr2* (Kamiya et al. 2012), a gene essential for testicular development, as natural mutants or knock-out of it leads to male-to-female sex reversal in several teleost species (Morinaga et al. 2007, Pan et al. 2022). The features of *amhr2* homologs that function as MSD genes in pufferfishes and catfishes differ from those observed in *P. flavescens*. In the Tiger pufferfish, sex determination hinges on a single amino-acid substitution distinguishing sex-specific *amhr2* alleles, with only the male allele capable of initiating testis development (Kamiya et al. 2012). In contrast, in the catfish family *amhr2* underwent a duplication creating a truncated copy that took the role of MSD gene (Wen et al. 2022). The *P. flavescens amhr2y* paralog resembles the catfish system, but the divergence of the new copy is tightly linked to the expansion of intronic satDNA. This peculiar architecture is reminiscent of “gigantic” genes in fruit files, where the presence of intronic satDNA influences the timing of gene expression during spermatogenesis (Fingerhut et al. 2019). Hence, a definitive identification of the MSD gene in *P. flavescen*s, combined with a characterization of its functional and evolutionary properties, will contribute to a broader understanding of vertebrate sex-determination mechanisms.

## Conclusion

By combining genomic investigations and demographic surveys, over a wide latitudinal range and across several life stages, we were able to narrow down the mechanisms driving Adult Sex Ratio (ASR) variation in natural populations of *P. flavescens*. While we cannot eliminate a small contribution of the SD system, we are confident that SD, genetic or environmental, cannot fully explain the ASR patterns observed. Breeding related mortality, although likely to occur in *P. flavescens*, is not a sufficient explanation either. Our favored hypothesis is sex-specific habitat distribution, which would explain the sporadic observation of male bias, and the absence of sex bias in pre-breeding adults. With this work, we have contributed important knowledge to the establishment of *P. flavescens* as a model system for the study of climatic effects on marine populations. Such model systems are going to be essential to understand the interacting roles of evolution, ecology, and demography in determining the response of natural populations to the ongoing climate change.

## Data availability

The newly-generated genomic data for this study have been deposited in the European Nucleotide Archive (ENA) at EMBL-EBI under accession number PRJEB103015 (https://www.ebi.ac.uk/ena/browser/view/PRJEB103015). The ddRAD reads have been deposited at NCBI, bioproject 1224570: http://www.ncbi.nlm.nih.gov/bioproject/1224570. Snakemake pipelines of most bioinformatic analyses are available at https://github.com/SLAment/GobySD.

## Field collections

For the collection of the 2022 field data, we extend our gratitude to Claudia Aparicio Estalella, Mathias Nyheim, Vera Løvold, Ida Lie-Nilsen, Dagmar M. Coelle, Paula Schmidtz, Henri Martinossi, Karoline Brudevoll Rognlien, Pauline Mesterson Pitter, Erik Amundsen, Amy Li, and Audun Aas Roseth. For the collection of the historical data, we extend our gratitude to Kai Lindström, Kenyon Mobley and Karen de Jong. We would like to thank Octavio M. Palacios-Gimenez for the advice on satDNA annotation.

## References

Altschul, S. F., Madden, T. L., Schäffer, A. A., Zhang, J., Zhang, Z., Miller, W., & Lipman, D. J. (1997). Gapped BLAST and PSI-BLAST : a new generation of protein database search programs. 25(17), 3389–3402.

Amundsen, T. (2018). Sex roles and sexual selection: Lessons from a dynamic model system. Current Zoology, 64(3), 363–392.

Ancona, S., Dénes, F. V., Krüger, O., Székely, T., & Beissinger, S. R. (2017). Estimating adult sex ratios in nature. Philosophical Transactions of the Royal Society B: Biological Sciences, 372(1729), 20160313.

Andersson, M. (1994). Sexual selection (Vol. 72). Princeton University Press.

Arnqvist, G., & Rowe, L. (2005). Sexual conflict (Vol. 27). Princeton university press.

Aydın, M., & Karadurmuş, U. (2022). Population Dynamics, Current Trends and Future Prospects of the Black Goby (Gobius niger) in the Eastern Part of the Black Sea (Turkiye). Aquatic Sciences and Engineering, 37(3), 142–150.

Bachtrog, D., Mank, J. E., Peichel, C. L., Kirkpatrick, M., Otto, S. P., Ashman, T.-L., Hahn, M. W., Kitano, J., Mayrose, I., Ming, R., & others. (2014). Sex determination: Why so many ways of doing it? PLoS Biology, 12(7), e1001899.

Bao, Z., & Eddy, S. R. (2002). Automated De Novo Identification of Repeat Sequence Families in Sequenced Genomes. Genome Research, 12(8), 1269–1276. 10.1101/gr.88502

Baroiller, J.-F., & d’Cotta, H. (2016). The reversible sex of gonochoristic fish: Insights and consequences. Sexual Development, 10(5–6), 242–266.

Bates, D., Mächler, M., Bolker, B., & Walker, S. (2015). Fitting Linear Mixed-Effects Models Using lme4. Journal of Statistical Software, 67(1), 1–48. 10.18637/jss.v067.i01

Benson, G. (1999). Tandem repeats finder: A program to analyze DNA sequences. Nucleic Acids Research, 27(2), 573–580. 10.1093/nar/27.2.573

Campbell, M. S., Law, M., Holt, C., Stein, J. C., Moghe, G. D., Hufnagel, D. E., Lei, J., Achawanantakun, R., Jiao, D., Lawrence, C. J., Ware, D., Shiu, S.-H., Childs, K. L., Sun, Y., Jiang, N., & Yandell, M. (2014). MAKER-P: A Tool Kit for the Rapid Creation, Management, and Quality Control of Plant Genome Annotations. Plant Physiology, 164(2), 513–524. 10.1104/pp.113.230144

Catchen, J., Hohenlohe, P. A., Bassham, S., Amores, A., & Cresko, W. A. (2013). Stacks: An analysis tool set for population genomics. Molecular Ecology, 22(11), 3124–3140.

Chang, C. C., Chow, C. C., Tellier, L. C., Vattikuti, S., Purcell, S. M., & Lee, J. J. (2015). Second-generation PLINK: rising to the challenge of larger and richer datasets. Gigascience, 4(1), s13742–015.

Cheng, H., Concepcion, G. T., Feng, X., Zhang, H., & Li, H. (2021). Haplotype-resolved de novo assembly using phased assembly graphs with hifiasm. Nature Methods, 18(2), 170–175. 10.1038/s41592-020-01056-5

Clarke, T. (1983). Sex ratios and sexual differences in size among mesopelagic fishes from the central Pacific Ocean. Marine Biology, 73, 203–209.

Danecek, P., Bonfield, J. K., Liddle, J., Marshall, J., Ohan, V., Pollard, M. O., Whitwham, A., Keane, T., McCarthy, S., Davies, R., & others. (2021). Twelve years of SAMtools and BCFtools. GigaScience 10, giab008.

De Coster, W., & Rademakers, R. (2023). NanoPack2: Population-scale evaluation of long-read sequencing data. Bioinformatics, 39(5), btad311.

de Jong, K., Forsgren, E., Sandvik, H., & Amundsen, T. (2012). Measuring mating competition correctly: Available evidence supports operational sex ratio theory. Behavioral Ecology, 23(6), 1170–1177.

Donald, P. F. (2007). Adult sex ratios in wild bird populations. Ibis, 149(4), 671–692.

Duffy, T. A., Hice, L. A., & Conover, D. O. (2015). Pattern and scale of geographic variation in environmental sex determination in the Atlantic silverside, Menidia menidia. Evolution, 69(8), 2187–2195.

Düsing, K. (1884). Die regulierung des geschlechtsverhältnisses bei der vermehrung der menschen, tiere und pflanzen. Gustav Fischer.

Eberhart-Phillips, L. J., Küpper, C., Carmona-Isunza, M. C., Vincze, O., Zefania, S., Cruz-López, M., Kosztolányi, A., Miller, T. E., Barta, Z., Cuthill, I. C., & others. (2018). Demographic causes of adult sex ratio variation and their consequences for parental cooperation. Nature Communications, 9(1), 1651.

Edwards, A. (2000). Carl Düsing (1884) on the regulation of the sex-ratio. Theoretical Population Biology, 58(3), 255–257.

Emlen, S. T., & Oring, L. W. (1977). Ecology, sexual selection, and the evolution of mating systems. Science, 197(4300), 215–223.

Feron, R. (2025). sgtr: Visualize population genomics analyses results. https://github.com/SexGenomicsToolkit/sgtr

Feron, R., Pan, Q., Wen, M., Imarazene, B., Jouanno, E., Anderson, J., Herpin, A., Journot, L., Parrinello, H., Klopp, C., & others. (2021). RADSex: A computational workflow to study sex determination using restriction site-associated DNA sequencing data. Molecular Ecology Resources, 21(5), 1715–1731.

Fingerhut, J. M., Moran, J. V., & Yamashita, Y. M. (2019). Satellite DNA-containing gigantic introns in a unique gene expression program during Drosophila spermatogenesis. PLOS Genetics, 15(5), e1008028. 10.1371/journal.pgen.1008028

Flynn, Jullien M, Hubley, R., Goubert, C., Rosen, J., Clark, A. G., Feschotte, C., & Smit, A. F. (2020). RepeatModeler2 for automated genomic discovery of transposable element families. Proceedings of the National Academy of Sciences, 117(17), 9451–9457.

Flynn, Jullien M., Hubley, R., Goubert, C., Rosen, J., Clark, A. G., Feschotte, C., & Smit, A. F. (2020). RepeatModeler2 for automated genomic discovery of transposable element families. Proceedings of the National Academy of Sciences, 117(17), 9451–9457. 10.1073/pnas.1921046117

Forsgren, E., Amundsen, T., Borg, Å. A., & Bjelvenmark, J. (2004). Unusually dynamic sex roles in a fish. Nature, 429(6991), 551–554.

Fosså, J. H. (1991). The ecology of the two-spot goby (Gobiusculus flavescens Fabricius): The potential for cod enhancement.

Fu, L., Niu, B., Zhu, Z., Wu, S., & Li, W. (2012). CD-HIT: Accelerated for clustering the next-generation sequencing data. Bioinformatics, 28(23), 3150–3152. 10.1093/bioinformatics/bts565

Galvão, T. B., Bertollo, L. A. C., & Molina, W. F. (2011). Chromosomal complements of some Atlantic Blennioidei and Gobioidei species (Perciformes). Comparative Cytogenetics, 5(4), 259.

Gibson, R., & Ezzi, I. (1978). The biology of a Scottish population of Fries’ goby, Lesueurigobius friesii. Journal of Fish Biology, 12(4), 371–389.

Gremme, G., Steinbiss, S., & Kurtz, S. (2013). GenomeTools: A Comprehensive Software Library for Efficient Processing of Structured Genome Annotations. IEEE/ACM Transactions on Computational Biology and Bioinformatics, 10(03), 645–656. 10.1109/TCBB.2013.68

Hackl, T., Ankenbrand, M., van Adrichem, B., Wilkins, D., & Haslinger, K. (2024). Gggenomes: Effective and versatile visualizations for comparative genomics. *arXiv Preprint arXiv:2411.13556*.

Honeycutt, J., Deck, C., Miller, S., Severance, M., Atkins, E., Luckenbach, J., Buckel, J., Daniels, H., Rice, J., Borski, R., & others. (2019). Warmer waters masculinize wild populations of a fish with temperature-dependent sex determination. Scientific Reports, 9(1), 6527.

Jaenike, J. (2001). Sex Chromosome Meiotic Drive. Annual Review of Ecology and Systematics, 32(1), 25–49. 10.1146/annurev.ecolsys.32.081501.113958

Jenner, L. P., Peska, V., Fulnečková, J., & Sýkorová, E. (2022). Telomeres and Their Neighbors. Genes, 13(9), 1663. 10.3390/genes13091663

Jones, P., Binns, D., Chang, H.-Y., Fraser, M., Li, W., McAnulla, C., McWilliam, H., Maslen, J., Mitchell, A., Nuka, G., Pesseat, S., Quinn, A. F., Sangrador-Vegas, A., Scheremetjew, M., Yong, S.-Y., Lopez, R., & Hunter, S. (2014). InterProScan 5: Genome-scale protein function classification. Bioinformatics, 30(9), 1236–1240. 10.1093/bioinformatics/btu031

Kamiya, T., Kai, W., Tasumi, S., Oka, A., Matsunaga, T., Mizuno, N., Fujita, M., Suetake, H., Suzuki, S., Hosoya, S., Tohari, S., Brenner, S., Miyadai, T., Venkatesh, B., Suzuki, Y., & Kikuchi, K. (2012). A Trans-Species Missense SNP in Amhr2 Is Associated with Sex Determination in the Tiger Pufferfish, Takifugu rubripes (Fugu). PLoS Genetics, 8(7), e1002798. 10.1371/journal.pgen.1002798

Kappeler, P. M., Benhaiem, S., Fichtel, C., Fromhage, L., Höner, O. P., Jennions, M. D., Kaiser, S., Krüger, O., Schneider, J. M., Tuni, C., & others. (2023). Sex roles and sex ratios in animals. Biological Reviews, 98(2), 462–480.

Kasapoğlu, N. (2016). Age, growth and mortality rates of discard species (Uranoscopus scaber, Neogobius melanostomus and Gobius niger) in the Black Sea. Ege Journal of Fisheries and Aquatic Sciences, 33(4), 397–403.

Katoh, K., Rozewicki, J., & Yamada, Kazunori D. (2019). MAFFT online service: Multiple sequence alignment, interactive sequence choice and visualization. Briefings in Bioinformatics, 20(4), 1160–1166.

Katoh, K., Rozewicki, J., & Yamada, Kazunori D. (2019). MAFFT online service: Multiple sequence alignment, interactive sequence choice and visualization. Briefings in Bioinformatics, 20(4), 1160–1166. 10.1093/bib/bbx108

Kendall, N. W., & Quinn, T. P. (2013). Size-selective fishing affects sex ratios and the opportunity for sexual selection in Alaskan sockeye salmon Oncorhynchus nerka. Oikos, 122(3), 411–420.

Klinkhardt, M. (1992). Chromosome structures of four Norwegian gobies (Gobiidae, Teleostei) and a hypothetical model of their karyo-evolution. Chromatin, 3(1), 169–183.

Korunes, K. L., & Samuk, K. (2021). pixy: Unbiased estimation of nucleotide diversity and divergence in the presence of missing data. Molecular Ecology Resources, 22(3), 1228–1229. 10.1111/1755-0998.13326

Kurtz, S., Phillippy, A., Delcher, A. L., Smoot, M., Shumway, M., Antonescu, C., & Salzberg, S. L. (2004). Versatile and open software for comparing large genomes. Genome Biology, 5(2), R12.

Leder, E. H., André, C., Le Moan, A., Töpel, M., Blomberg, A., Havenhand, J. N., Lindström, K., Volckaert, F. A., Kvarnemo, C., Johannesson, K., & others. (2021). Post-glacial establishment of locally adapted fish populations over a steep salinity gradient. Journal of Evolutionary Biology, 34(1), 138–156.

Lema, S. C., Luckenbach, J. A., Yamamoto, Y., & Housh, M. J. (2024). Fish reproduction in a warming world: Vulnerable points in hormone regulation from sex determination to spawning. Philosophical Transactions of the Royal Society B, 379(1898), 20220516.

Li, H. (2018). Minimap2: Pairwise alignment for nucleotide sequences. Bioinformatics, 34(18), 3094–3100.

Li, H., & Durbin, R. (2009). Fast and accurate short read alignment with Burrows–Wheeler transform. Bioinformatics, 25(14), 1754–1760.

Magnhagen, C. (1991). Predation risk as a cost of reproduction. Trends in Ecology & Evolution, 6(6), 183–186.

Manni, M., Berkeley, M. R., Seppey, M., Simão, F. A., & Zdobnov, E. M. (2021). BUSCO update: Novel and streamlined workflows along with broader and deeper phylogenetic coverage for scoring of eukaryotic, prokaryotic, and viral genomes. Molecular Biology and Evolution, 38(10), 4647–4654.

Marçais, K., & Kingsford, C. (2011). JELLYFISH–fast, parallel k-mer counting for DNA. Bioinformatics, 27(6), 764–770.

Martinossi-Allibert, I., Ratikainen, I., Albretsen, J., Eknes, E. J., Ellingsen, I., Estalella, C. A., Jensen, H., Raeymaekers, J., Røilid, T., Sodeland, M., & others. (2025). Meeting the challenge of latitude: A climatic gradient shapes the reproductive strategy of a nest-brooding marine fish. Marine Ecology Progress Series, 767, 101–120.

Martinossi-Allibert, I., Wacker, S., Aparicio Estalella, C., Kvarnemo, C., & Amundsen, T. (2025). A test of operational sex ratio theory across latitudes reveals temporal variation in sex-specific behavioural reaction norms. Journal of Animal Ecology, 94(4), 642–656.

McKenna, A., Hanna, M., Banks, E., Sivachenko, A., Cibulskis, K., Kernytsky, A., Garimella, K., Altshuler, D., Gabriel, S., Daly, M., & DePristo, M. A. (2010). The genome analysis toolkit: A MapReduce framework for analyzing next-generation DNA sequencing data. Genome Research, 20(9), 1297–1303. 10.1101/gr.107524.110

Minh, B. Q., Schmidt, H. A., Chernomor, O., Schrempf, D., Woodhams, M. D., Von Haeseler, A., & Lanfear, R. (2020). IQ-TREE 2: New models and efficient methods for phylogenetic inference in the genomic era. Molecular Biology and Evolution, 37(5), 1530–1534.

Miyoshi, K., Hattori, R. S., Strüssmann, C. A., Yokota, M., & Yamamoto, Y. (2020). Phenotypic/genotypic sex mismatches and temperature-dependent sex determination in a wild population of an old world atherinid, the cobaltcap silverside Hypoatherina tsurugae. Molecular Ecology, 29(13), 2349–2358.

Morinaga, C., Saito, D., Nakamura, S., Sasaki, T., Asakawa, S., Shimizu, N., Mitani, H., Furutani-Seiki, M., Tanaka, M., & Kondoh, H. (2007). The hotei mutation of medaka in the anti-Müllerian hormone receptor causes the dysregulation of germ cell and sexual development. Proceedings of the National Academy of Sciences, 104(23), 9691–9696.

Myhre, L. C., de Jong, K., Forsgren, E., & Amundsen, T. (2012). Sex roles and mutual mate choice matter during mate sampling. The American Naturalist, 179(6), 741–755.

Nguyen, N. T. T., Vincens, P., Dufayard, J. F., Roest Crollius, H., & Louis, A. (2022). Genomicus in 2022: Comparative tools for thousands of genomes and reconstructed ancestors. Nucleic Acids Research, 50(D1), D1025–D1031. 10.1093/nar/gkab1091

Novak, P. (2025, November). TideCluster. Zenodo. 10.5281/zenodo.17638302

Oliveira, M., Costa, E., Araújo, A., Pessoa, E., Carvalho, M., Cavalcante, L., & Chellappa, S. (2012). Sex ratio and length-weight relationship for five marine fish species from Brazil. Journal of Marine Biology & Oceanography, 1, 2.

Ospina-Alvarez, N., & Piferrer, F. (2008). Temperature-dependent sex determination in fish revisited: Prevalence, a single sex ratio response pattern, and possible effects of climate change. PloS One, 3(7), e2837.

Ou, S., & Jiang, N. (2018). LTR_retriever: A Highly Accurate and Sensitive Program for Identification of Long Terminal Repeat Retrotransposons. Plant Physiology, 176(2), 1410–1422. 10.1104/pp.17.01310

Pan, Q., Herpin, A., & Guiguen, Y. (2022). Inactivation of the anti-Müllerian hormone receptor type 2 (amhrII) gene in Northern Pike (Esox lucius) results in male-to-female sex reversal. Sexual Development, 16(4), 289–294.

Pan, Q., Kay, T., Depincé, A., Adolfi, M., Schartl, M., Guiguen, Y., & Herpin, A. (2021). Evolution of master sex determiners: TGF-β signalling pathways at regulatory crossroads. Philosophical Transactions of the Royal Society B, 376(1832), 20200091. 10.1098/rstb.2020.0091

Prazdnikov, D. V. (2023). Chromosome complement of Pomatoschistus marmoratus and Karyotype evolution in sand gobies (Gobiidae: Gobionellinae). Turkish Journal of Fisheries and Aquatic Sciences, 23(7).

Price, A. L., Jones, N. C., & Pevzner, P. A. (2005). De novo identification of repeat families in large genomes. Bioinformatics, 21(suppl_1), i351–i358. 10.1093/bioinformatics/bti1018

Purcell, S., Neale, B., Todd-Brown, K., Thomas, L., Ferreira, M. A., Bender, D., Maller, J., Sklar, P., De Bakker, P. I., Daly, M. J., & others. (2007). PLINK: a tool set for whole-genome association and population-based linkage analyses. The American Journal of Human Genetics, 81(3), 559–575.

Quinlan, A. R., & Hall, I. M. (2010). BEDTools: A flexible suite of utilities for comparing genomic features. Bioinformatics, 26(6), 841–842.

R Core Team. (2025). R: A Language and Environment for Statistical Computing. R Foundation for Statistical Computing. https://www.R-project.org/

Rundio, D. E., Williams, T. H., Pearse, D. E., & Lindley, S. T. (2012). Male-biased sex ratio of nonanadromous Oncorhynchus mykiss in a partially migratory population in California. Ecology of Freshwater Fish, 21(2), 293–299.

Sadovy, Y., & Colin, P. (1995). Sexual development and sexuality in the Nassau grouper. Journal of Fish Biology, 46(6), 961–976.

Seger, J., & Stubblefield, J. W. (2002). Models of sex ratio evolution. Sex Ratios: Concepts and Research Methods, 2–25.

Skolbekken, R., & Utne-Palm, A. C. (2001). Parental investment of male two-spotted goby, Gobiusculus flavescens (Fabricius). Journal of Experimental Marine Biology and Ecology, 261(2), 137–157.

Smit, A., & Hubley, R. (2025, March). RepeatMasker. https://github.com/Dfam-consortium/RepeatMasker

Stanke, M., Keller, O., Gunduz, I., Hayes, A., Waack, S., & Morgenstern, B. (2006). AUGUSTUS: ab initio prediction of alternative transcripts. Nucleic Acids Research, 34(Web Server), W435–W439. 10.1093/nar/gkl200

Székely, T., Weissing, F. J., & Komdeur, J. (2014). Adult sex ratio variation: Implications for breeding system evolution. Journal of Evolutionary Biology, 27(8), 1500–1512.

Thompson, A. W., Hawkins, M. B., Parey, E., Wcisel, D. J., Ota, T., Kawasaki, K., Funk, E., Losilla, M., Fitch, O. E., Pan, Q., Feron, R., Louis, A., Montfort, J., Milhes, M., Racicot, B. L., Childs, K. L., Fontenot, Q., Ferrara, A., David, S. R., … Braasch, I. (2021). The bowfin genome illuminates the developmental evolution of ray-finned fishes. Nature Genetics, 53(9), 1373–1384. 10.1038/s41588-021-00914-y

Trochet, A., Courtois, E. A., Stevens, V. M., Baguette, M., Chaine, A., Schmeller, D. S., Clobert, J., & Wiens, J. J. (2016). Evolution of sex-biased dispersal. The Quarterly Review of Biology, 91(3), 297–320.

Uliano-Silva, M., Ferreira, J. G. R., Krasheninnikova, K., Formenti, G., Abueg, L., Torrance, J., Myers, E. W., Durbin, R., Blaxter, M., & others. (2023). MitoHiFi: A python pipeline for mitochondrial genome assembly from PacBio high fidelity reads. BMC Bioinformatics, 24(1), 288.

Untergasser, A., Cutcutache, I., Koressaar, T., Ye, J., Faircloth, B. C., Remm, M., & Rozen, S. G. (2012). Primer3—New capabilities and interfaces. Nucleic Acids Research, 40(15), e115–e115.

Utne-Palm, A. C., Eduard, K., Jensen, K. H., Mayer, I., & Jakobsen, P. J. (2015). Size dependent male reproductive tactic in the two-spotted goby (Gobiusculus flavescens). PLoS One, 10(12), e0143487.

Valdivieso, A., Wilson, C. A., Amores, A., da Silva Rodrigues, M., Nóbrega, R. H., Ribas, L., Postlethwait, J. H., & Piferrer, F. (2022). Environmentally-induced sex reversal in fish with chromosomal vs. Polygenic sex determination. Environmental Research, 213, 113549.

Wacker, S., Amundsen, T., Forsgren, E., & Mobley, K. B. (2014). Within-season variation in sexual selection in a fish with dynamic sex roles. Molecular Ecology, 23(14), 3587–3599.

Wen, M., Pan, Q., Jouanno, E., Montfort, J., Zahm, M., Cabau, C., Klopp, C., Iampietro, C., Roques, C., Bouchez, O., Castinel, A., Donnadieu, C., Parrinello, H., Poncet, C., Belmonte, E., Gautier, V., Avarre, J., Dugue, R., Gustiano, R., … Guiguen, Y. (2022a). An ancient truncated duplication of the anti-Müllerian hormone receptor type 2 gene is a potential conserved master sex determinant in the Pangasiidae catfish family. Molecular Ecology Resources, 22(6), 2411–2428. 10.1111/1755-0998.13620

Wen, M., Pan, Q., Jouanno, E., Montfort, J., Zahm, M., Cabau, C., Klopp, C., Iampietro, C., Roques, C., Bouchez, O., Castinel, A., Donnadieu, C., Parrinello, H., Poncet, C., Belmonte, E., Gautier, V., Avarre, J., Dugue, R., Gustiano, R., … Guiguen, Y. (2022b). An ancient truncated duplication of the anti-Müllerian hormone receptor type 2 gene is a potential conserved master sex determinant in the Pangasiidae catfish family. Molecular Ecology Resources, 22(6), 2411–2428. 10.1111/1755-0998.13620

Wheeler, T. J. (2009). Large-scale neighbor-joining with NINJA. In S. L. Salzberg & T. Warnow (Eds.), Proceedings of the 9th Workshop on Algorithms in Bioinformatics (WABI) (pp. 375–389). Springer.

