## Supplementary figures and tables for "Female-biased sex ratios despite stable genetic sex determination across a climatic gradient in a marine fish"

*shared last-authorship

^1^Department of Zoology, Stockholm University, 106 91 Stockholm, Sweden

^2^Department of Biology, Norwegian University of Science and Technology (NTNU), Høgskoleringen 5, 7034 Trondheim, Norway

^3^Norwegian Institute for Nature Research (NINA), Høgskoleringen 9, 7034 Trondheim, Norway

^4^Institute of Marine Research, Nye Flødevigveien 20, 4817 His, Norway

^5^Centre for Coastal Research, Department of Natural Sciences, University of Agder, 4630 Kristiansand, Norway

^6^ Department of Immunology, Genetics and Pathology, Uppsala Genome Center, National Genomics Infrastructure Hosted by SciLifeLab, Uppsala University, Uppsala, Sweden

^7^Fish Capture Division, Institute of Marine Research (IMR), Nordnesgaten 50, 5005 Bergen, Norway

^8^Max Planck Institute for Biology, 72076, Tübingen, Germany

^9^Faculty of Biosciences and Aquaculture, Nord University, Bodø, Norway

**Table S1.** **Location of the markers used for sexing based on the *P. minutus* reference genome, along with genotypes in males and females of *P. flavescens*, and their corresponding primers.**

| ***P. minutus* Scaffold** | **SNP Position** | **Female gen.** | **Male gen.** | **Final sexing marker** | **Forward**  **primer (5'-3')** | **Reverse**  **primer (5'-3')** |
| --- | --- | --- | --- | --- | --- | --- |
| scaffold709 | 69411 | A/A | A/T | Discarded | TGCAGCCAGGCTAATGGAT | CCATTAGTCTTCATGAAATAGGTAAC |
| scaffold709 | 69413 | G/G | G/T | Discarded | TGCAGCCAGGCTAATGGAT | CCATTAGTCTTCATGAAATAGGTAAC |
| scaffold709 | 74649 | C/C | C/T | Kept | TGCAGGCCTAAACAGTGTCG | CGCTGCGTGACATGAACTC |
| scaffold709 | 104060 | A/A | A/C | Kept | TCATATGTGGGGGAATTACAGCT | GCAGTGTAGTTTCCCCTCCC |
| scaffold2702 | 47682 | T/T | T/A | Discarded | TGCAGGGATCTCTACACACG | TCCCCTCCTGCACATTTTGT |
| scaffold3260 | 39920 | G/G | G/A | Kept | CAGTGTGTGCTCAACTGAGAATT | TCGACTGGACTGACTCTTCA |
| scaffold3962 | 25171 | C/C | C/T | Kept | TGCAGTCCAACACACAGCC | CACACAGAAACGCACGCAC |
| scaffold4193 | 6290 | A/A | A/G | Discarded | TGCAGCTTTGTCTATGAGGTCA | TGTGATCCTGTTCTTTGAAAGGC |
| scaffold7908 | 19077 | C/C | C/T | Kept | TGCAGTACAGTGTGCATGGT | TGCTCTAGTCCAAAGTAAAATGCT |

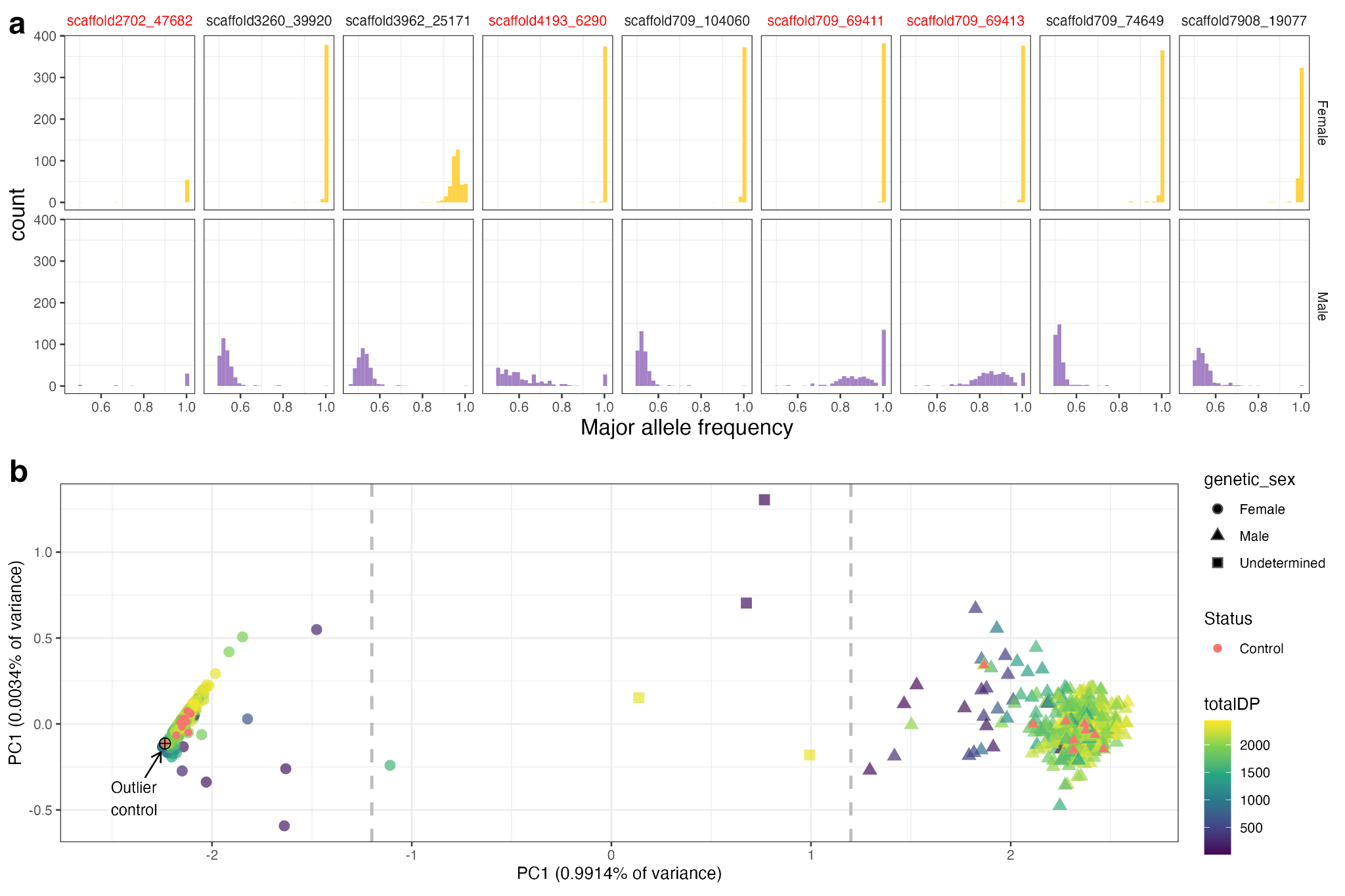

**Figure S1. Design of sex-specific markers for the two-spotted goby.** *In (****a****) the plot depicts the major allele frequency (MAF) distribution in the read counts of each marker for all eggs, larvae, and adults of unknown sex.* *Nine markers were selected from ddRAD polymorphisms mapped to the P. minutus reference genome (before the production of P. flavescens assemblies) associated to sex.* *Based on their MAF, the markers labeled in red were discarded, leaving five final markers for sexing. A given genotyped individual was considered genetically female if the average MAF of all five remaining markers was >0.85, male if <0.65, and otherwise undetermined. (****b****) A principal component analysis of the allele frequencies of surviving markers confirms a good separation between males and females, although a few samples are relatively intermediate (framed by dashed lines between -1.2 and 1.2 on the PC1 axis). Each point represents an individual. Control represents adults sexed based on phenotype. One phenotypically male control resulted in a genetic female (marked as an outlier). totalDP: sum of the allele depth of all five markers per individual.*

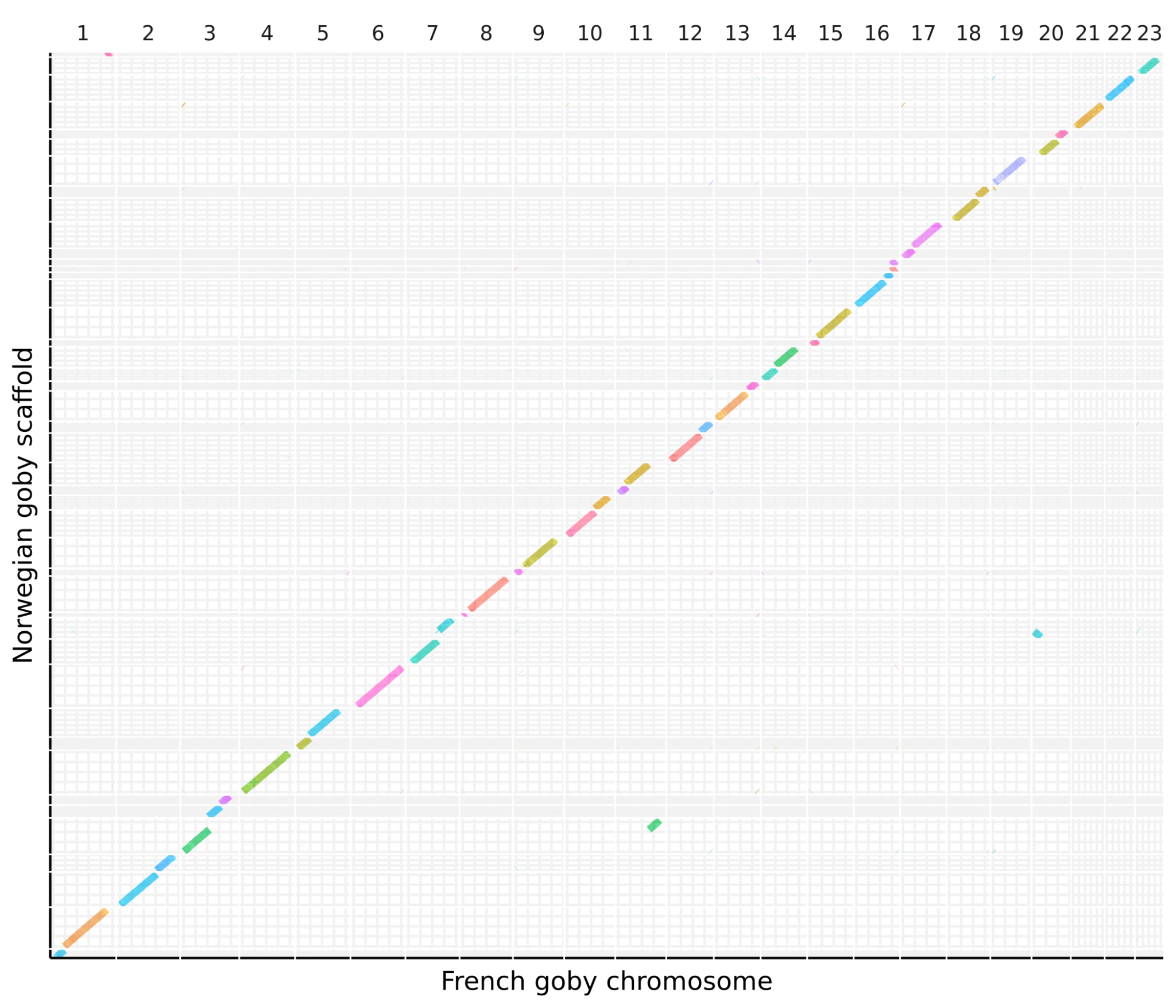

**Figure S2. Dot plot between the reference genome assembly of the France individual (fGobFla1) and the primary assembly of the Norwegian TH1 individual.** *Colors correspond to the different TH1 scaffolds. Only alignments larger than 30 Kbp with identity higher than 96% were considered. TH1 scaffolds smaller than 3 Mbp were ignored for clarity.*

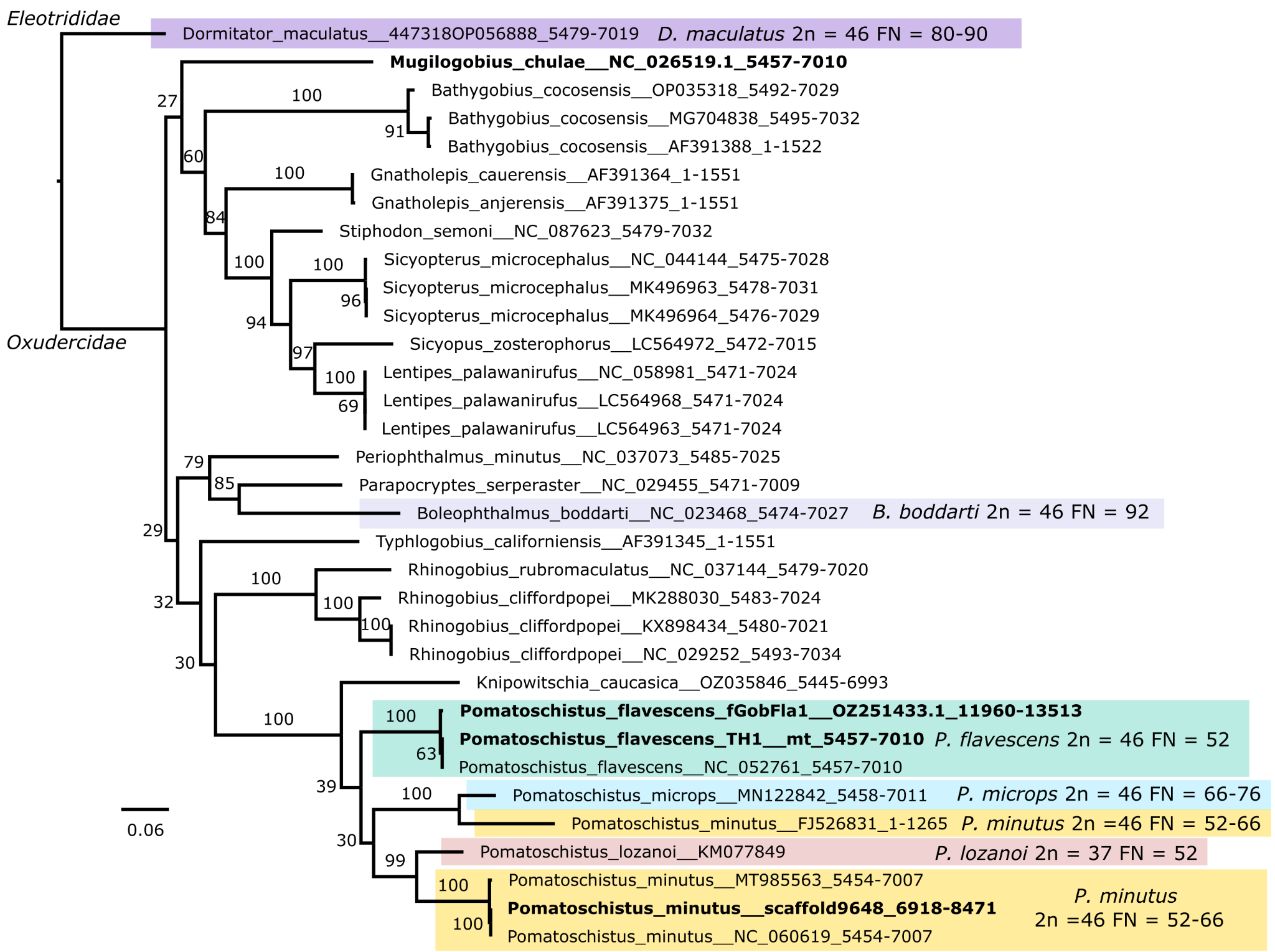

**Figure S3. Maximum likelihood phylogeny of the *cox1* gene of *P. flavescens* and closely related goby species.** *Species with known karyotypes are highlighted in colored boxes. Sequences obtained from whole-genome assemblies are highlighted in bold, with the exception of Mugilogobius chulae for which the mitogenome likely belongs to a different individual than that of the nuclear assembly. Branch support values correspond to non-parametric bootstraps. Branch lengths are drawn proportional to the scale bar (nucleotide substitutions per site).*

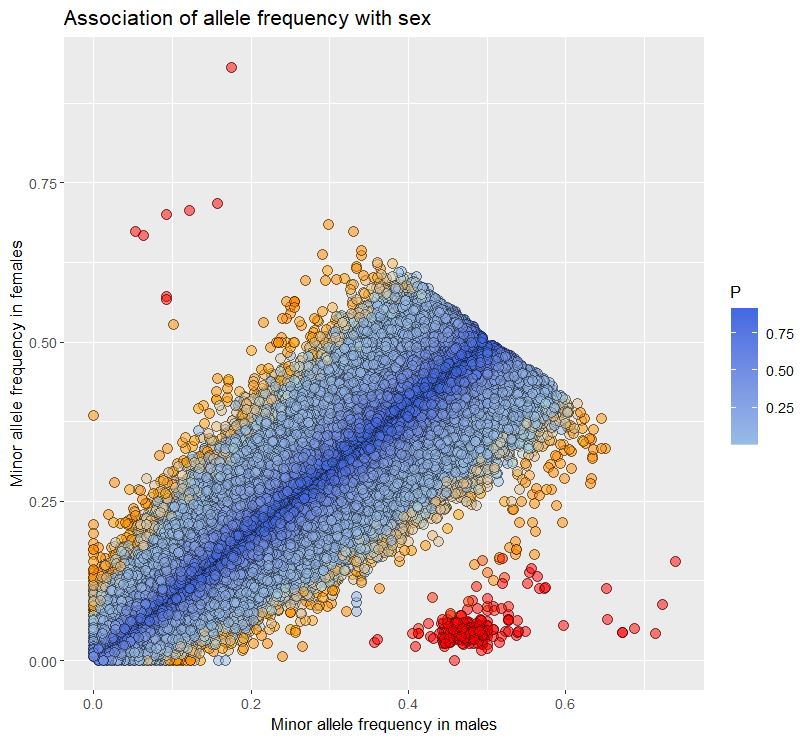

**Figure S4. Sex-specific allele frequency of ddRAD Single Nucleotide Polymorphisms (SNPs) shared among males and females.** Each dot is a SNP present in males and females, with the minor allele frequency in males on the x-axis and in females on the y-axis. The colour of the dots represents the p-value of a Fisher exact test of association between the allele frequency and phenotypic sex (p value: 1> Dark blue >10^-2^> Light blue >10^-3^> Orange >10^-10^> Red). Candidate SNPs for the sex determination locus are the cloud of red dots at the bottom right of the plot, close to intermediated frequency in males and close to fixation in females.

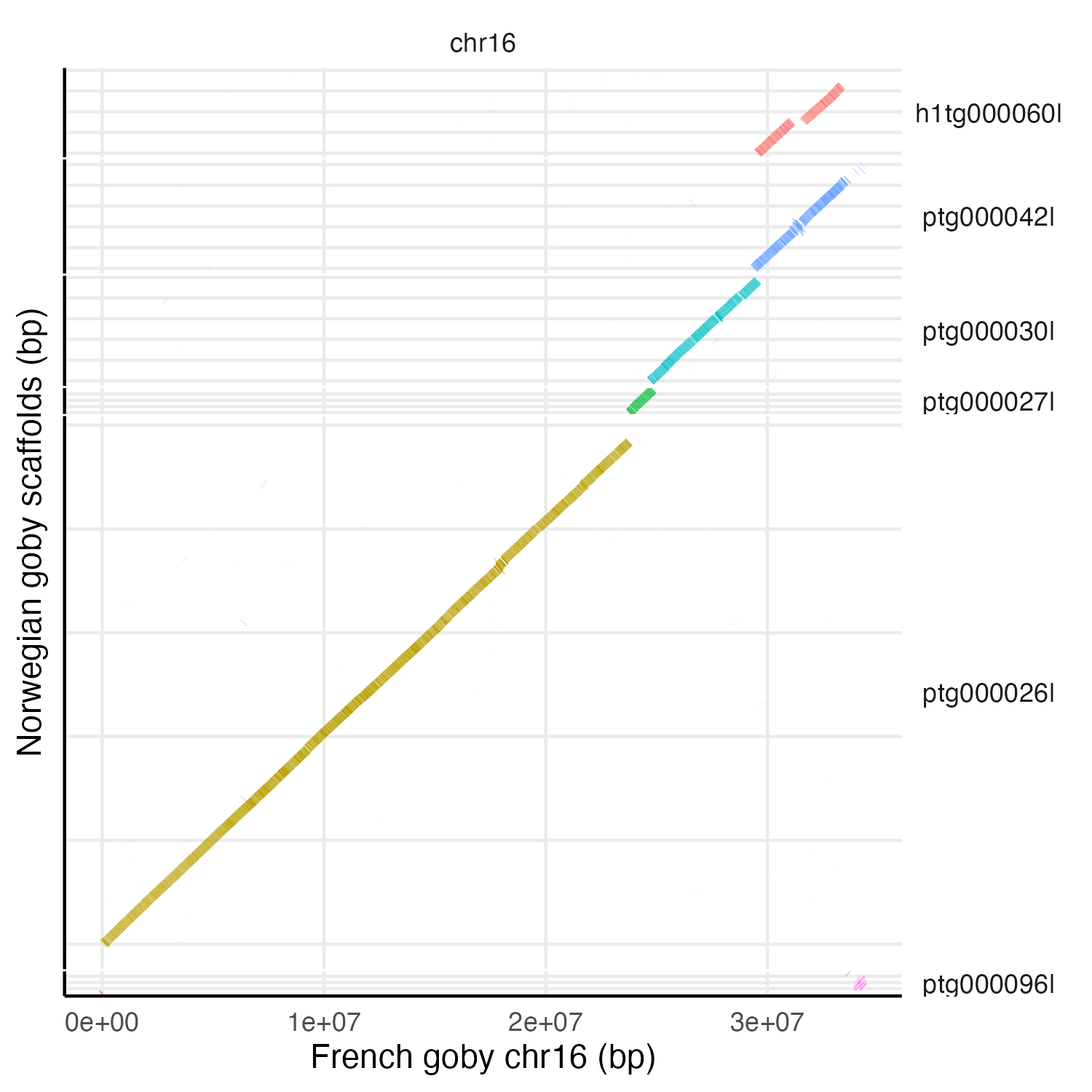

**Figure S5. Dot plot between the chromosome 16 of the French individual (fGobFla1) and the homologous scaffolds in the Norwegian individual (TH1) assembly.** *Scaffolds that start with “ptg” belong in the primary assembly, whilte the SD scaffold h1th000060l corresponds to the alternative haplotype of ptg000042l (= h2tg000053l). Colors correspond to the different TH1 scaffolds. Only alignments larger than 100 Kbp with identity higher than 96% were considered. Based on similarity, the chromosome 16 of fGobFla1 corresponds to the haplotype 2 of TH1.*

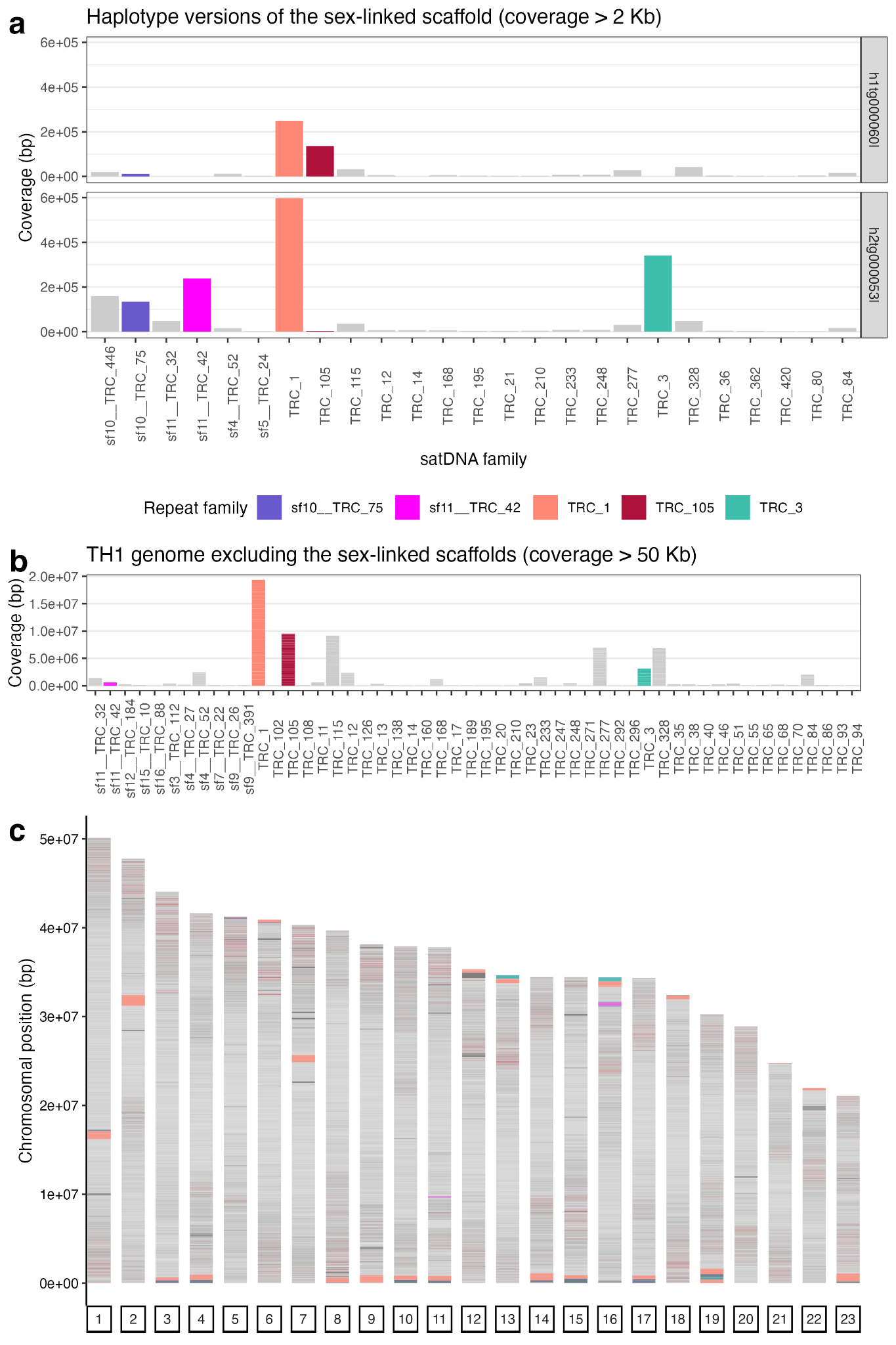

**Figure S6. Abundance of TideCluster satDNA families in the *P. flavescens* genome.** *In (****a****) the abundance in the two haplotype versions of the sex-linked scaffold is shown for families with an overall coverage larger than 2 Kb (i.e. uncommon families are ignored). In* ***(b),*** *the abundance of satDNA families across the TH1 genome but excluding the sex-linked scaffolds is shown for families with a coverage larger than 50 Kb. In (****c****) the distribution of satDNA families is displayed along the chromosomes of the fGobFla1 reference. The satDNA families that flank the SD region in chromosome 16 are highlighted with different colors.*

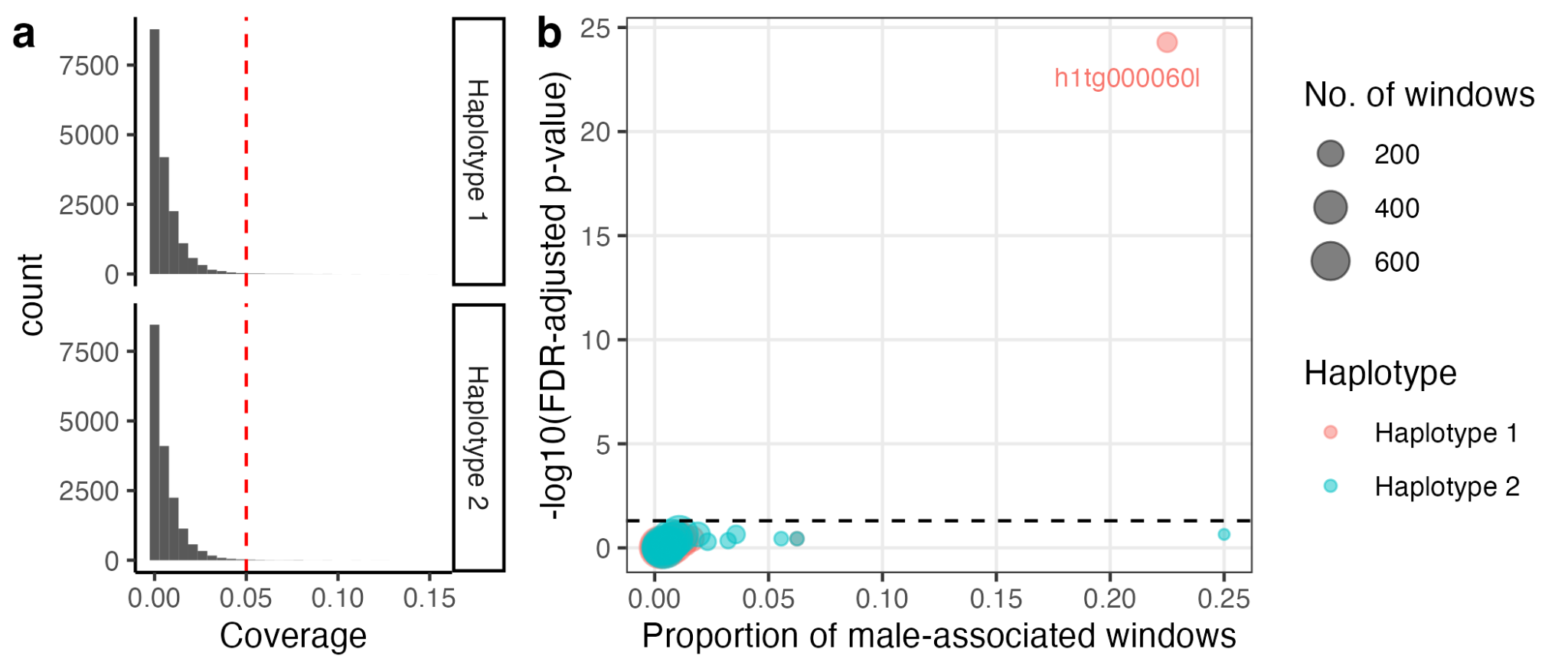

**Figure S7. Enrichment of male-associated k-mers in the TH1 haplotype-resolved assemblies.** *Coverage corresponds to the fraction of 50Kb-long non-overlapping windows that contain male-associated k-mers. In both haplotype-resolved assemblies, most windows have very low coverage (****a****). Setting 0.05 as an arbitrary threshold of coverage makes evident that a single scaffold in the haplotype 1 assembly is signficantly enriched for male-associated kmers (****b****). The black doted line corresponds to the significance level -log10(α), where α = 0.05. FDR: Benjamini–Hochberg false discovery rate.*

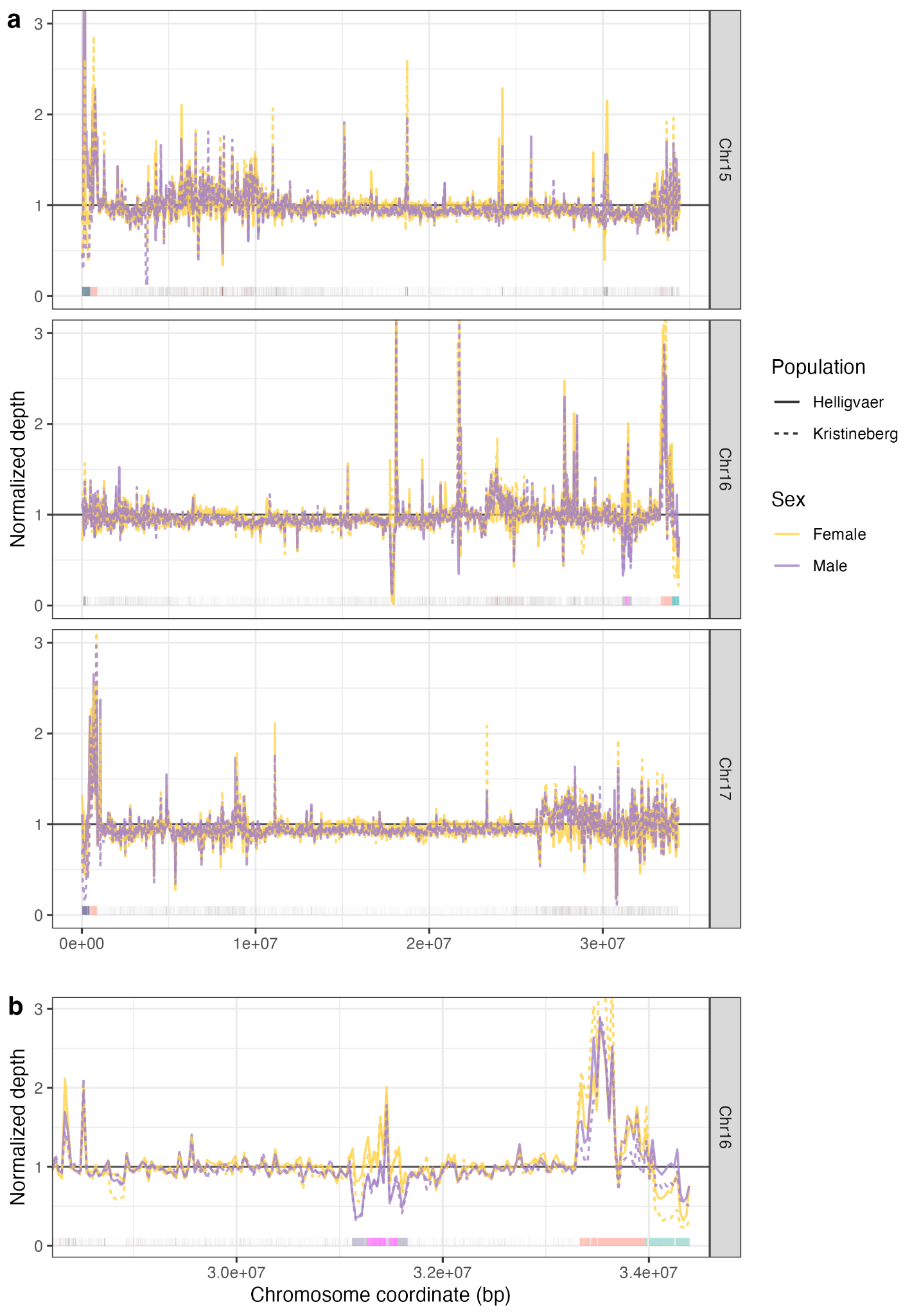

**Figure S8. Normalized coverage of four individuals sequenced with Illumina along selected chromosomes of fGobFla1.** *Depth of coverage was calculated in non-overlapping 30 Kb-long windows and normalized per individual using its whole-genome average. Normalized coverage is congruent between sexes (****a****), including the area corresponding to the sex-linked region in chromosome 16, except for the satDNA areas (****b****). The satDNA families that flank the sex-linked region in chromosome 16 are highlighted with different colors. Maximum normalized coverage was caped at 3 for clarity.*

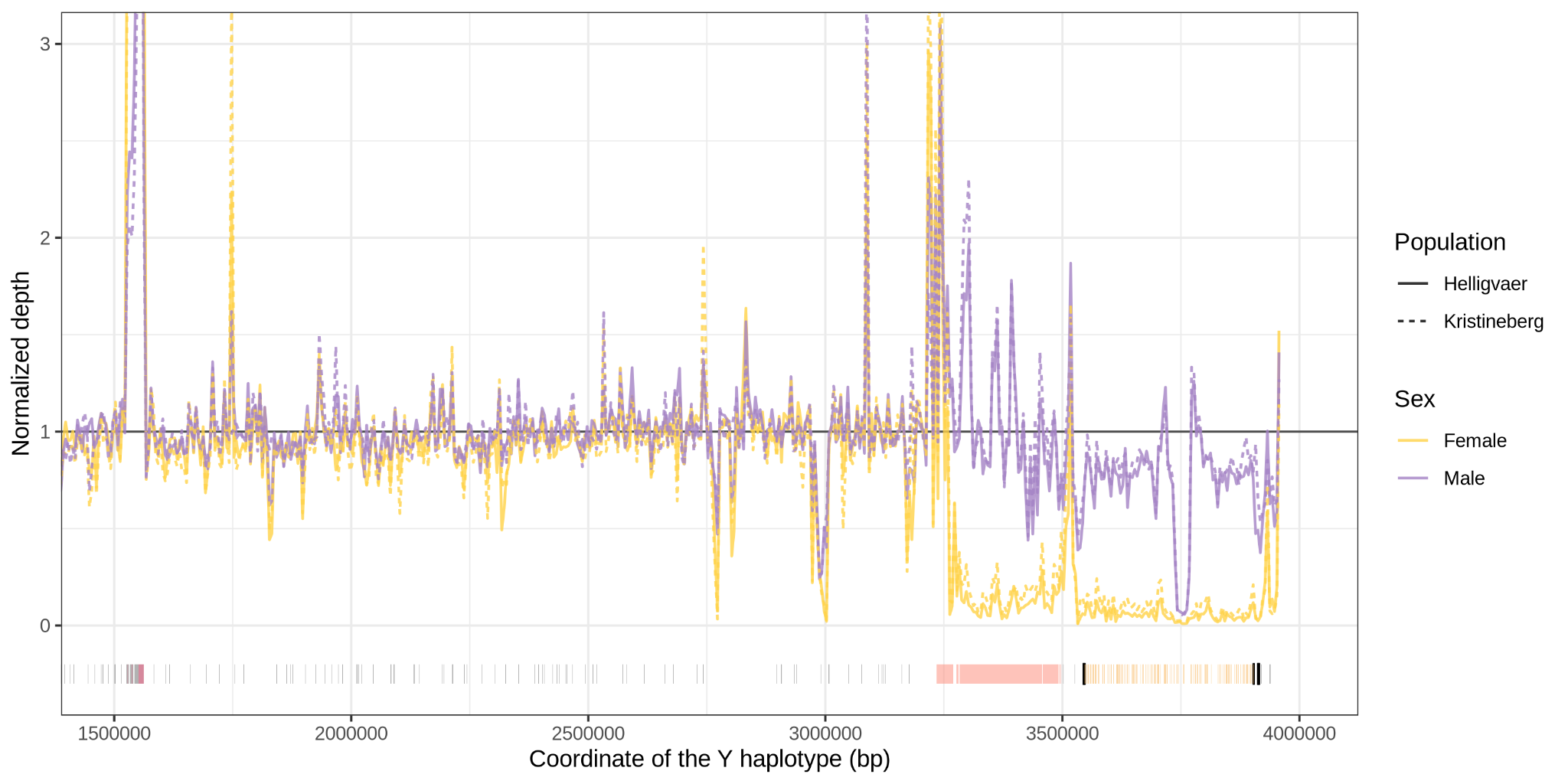

**Figure S9. Normalized coverage of four individuals sequenced with Illumina along the scaffold containing the SD region (Y haplotype).** *Depth of coverage was calculated in non-overlapping 5 Kb-long windows and normalized per individual using its whole-genome average. The satDNA families are highlighted with different colors at the bottom, while the exons of* amhr2y *are marked in black. Maximum normalized coverage was caped at 3 for clarity.*

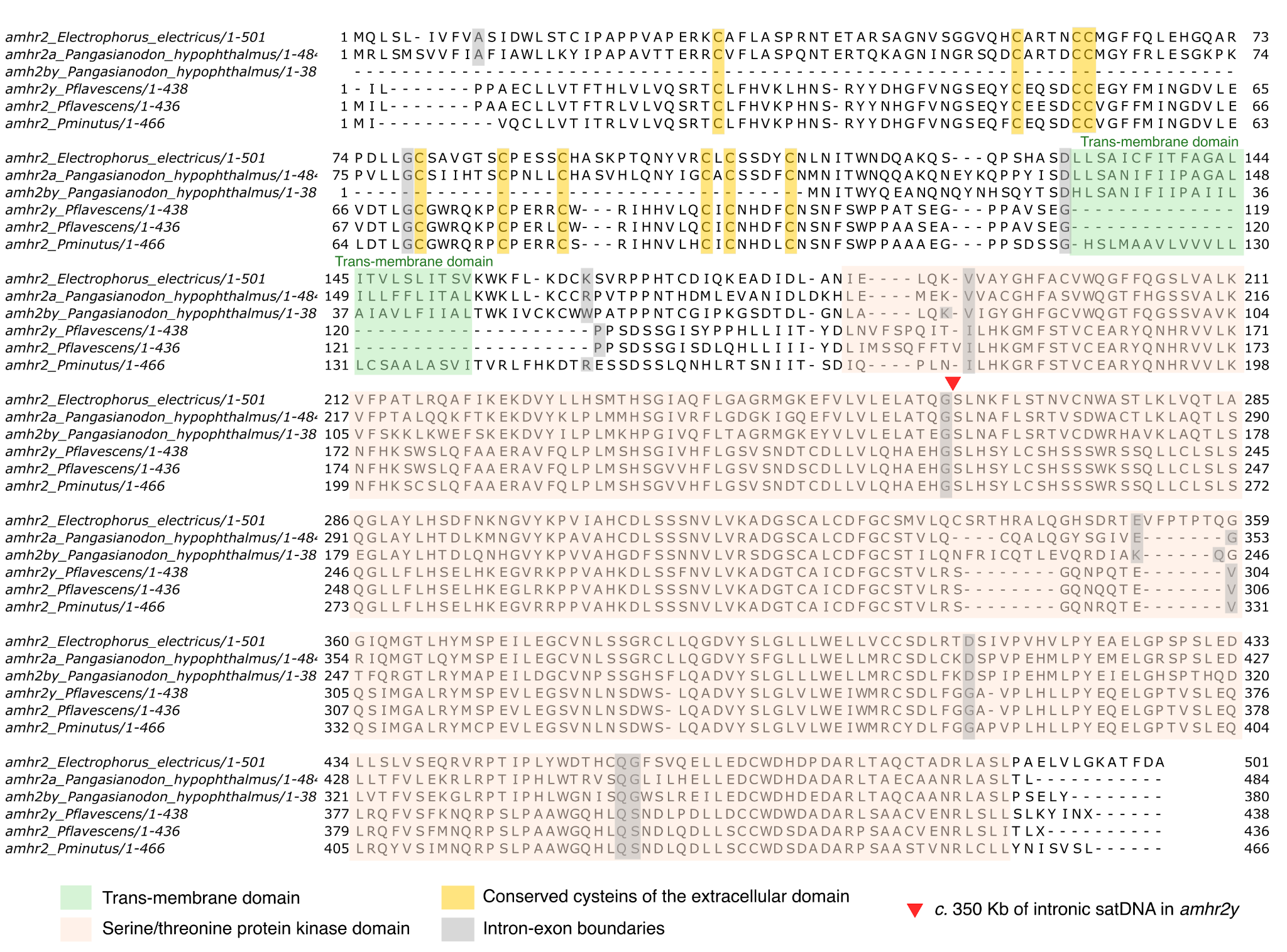

**Figure S10. Protein alignment of the *P. minutus amhr2* genes and homologs from other species.** *The homolog of* P. minutus *was obtained from transcriptomic data. The other three sequences represent the two* amhr2 *copies present in the* Pangasiidae *catfish family (from* Pangasianodon hypophthalmus*), as well as the ortholog in the electric eel (*Electrophorus electricus*) from the* Gymnotidae *family. As described in Wen et al. (2022), the duplication of* amhr2 *that acts as a MSD gene in the* Pangasiidae *fish (*amhr2by*) lost its N-terminus, corresponding to the extracellular domain. The two copies present in* P. flavescens *have the N-terminus, but the one in the SD region (*amhr2y*) has putatively lost its start codon, raising the possibility of a secondary start codon downstream (not to mention the transcriptional challenge of splicing out nearly 350 Kb of intronic satDNA). Notice that the absence of a trans-membrane domain in* P. flavescens *might be due to limitations in the manual definition of the gene model rather than true absence. The highlighted protein features follow Wen et al. (2022).*

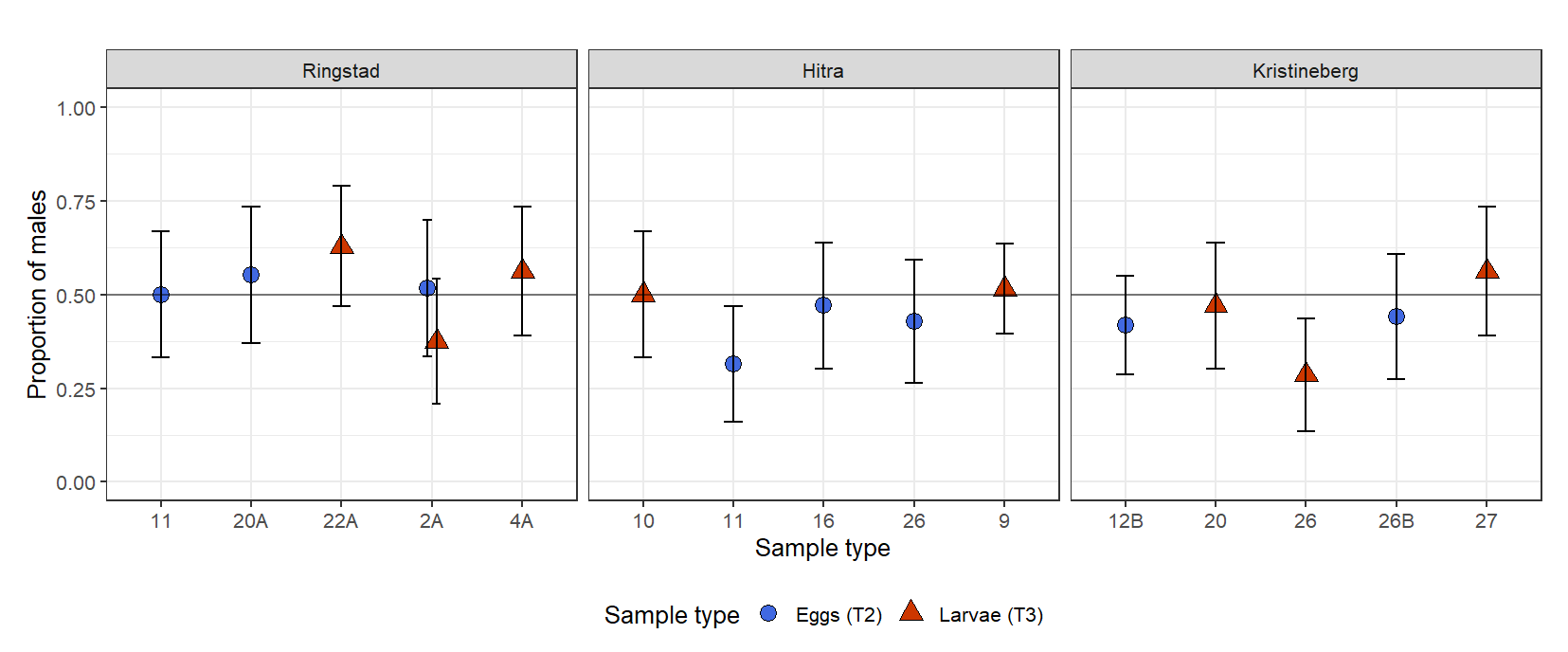

**Figure S11. Genotypic sex ratio of *P. flavescens* eggs and larvae in each clutch or nest, from three populations of the 2022 field season.** *The proportion of males identified by genotyping is given for each clutch or nest. Error bars represent a 95% confidence interval calculated according to the Normal approximation method. The range of sample size for each nest or clutch is between 29 and 66 individuals.*

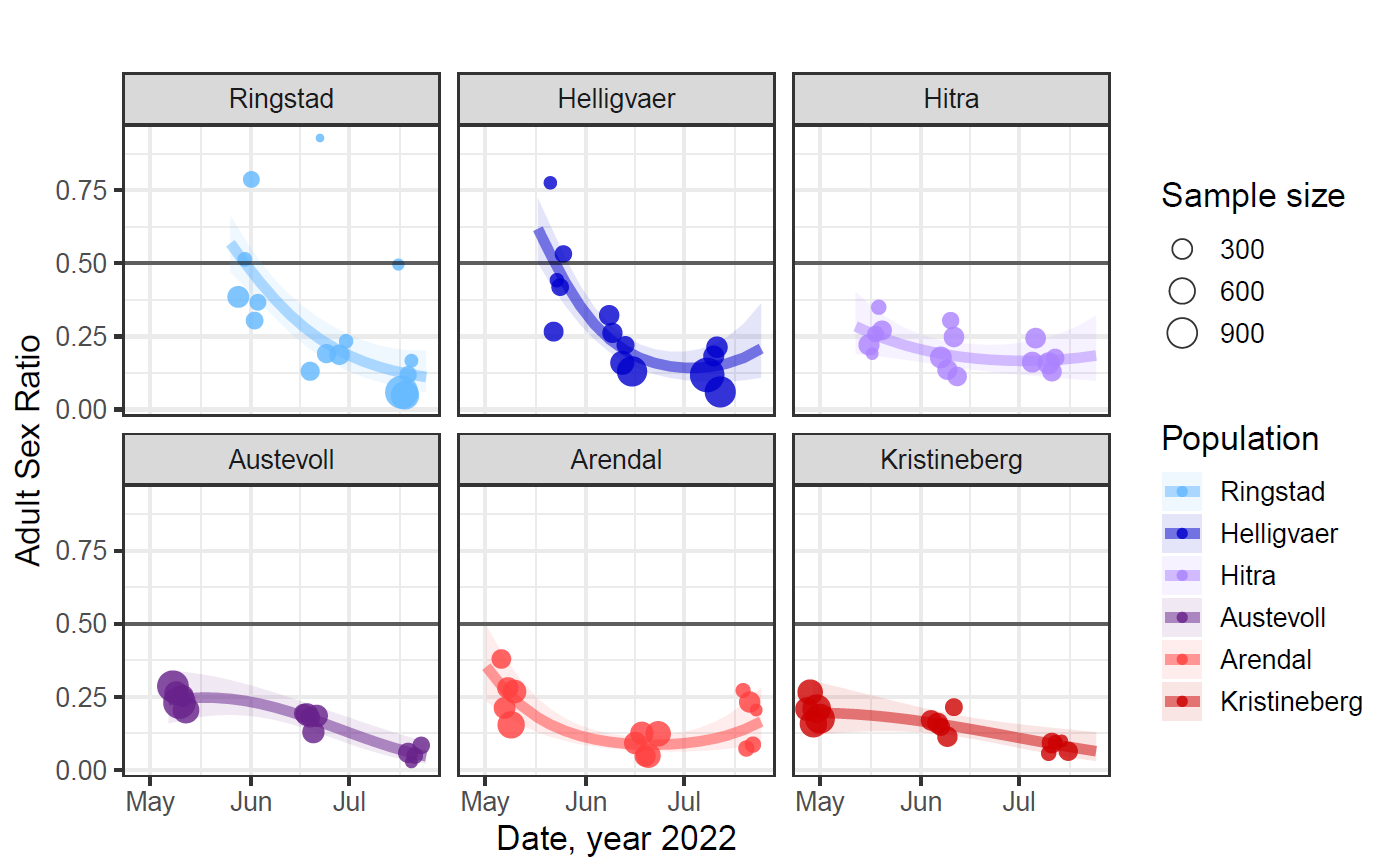

**Figure S12. Adult sex ratio (ASR) from observation transects during the 2022 breeding season, with one panel for each of the six Scandinavian populations.**

*Individuals were counted and sexed by snorkeling observers along transects. Each dot is a sampling transect, five per population and per time period (early, mid- and late- season). ASR is calculated as the proportion of males: m/(m+f). The size of each dot is proportional to the total adult census (m+f). The curves for each population represent the fit from a binomial model (see Table 1), with 95% confidence intervals encompassed by the shaded area around each curve. The horizontal black line indicates a balanced sex ratio.*

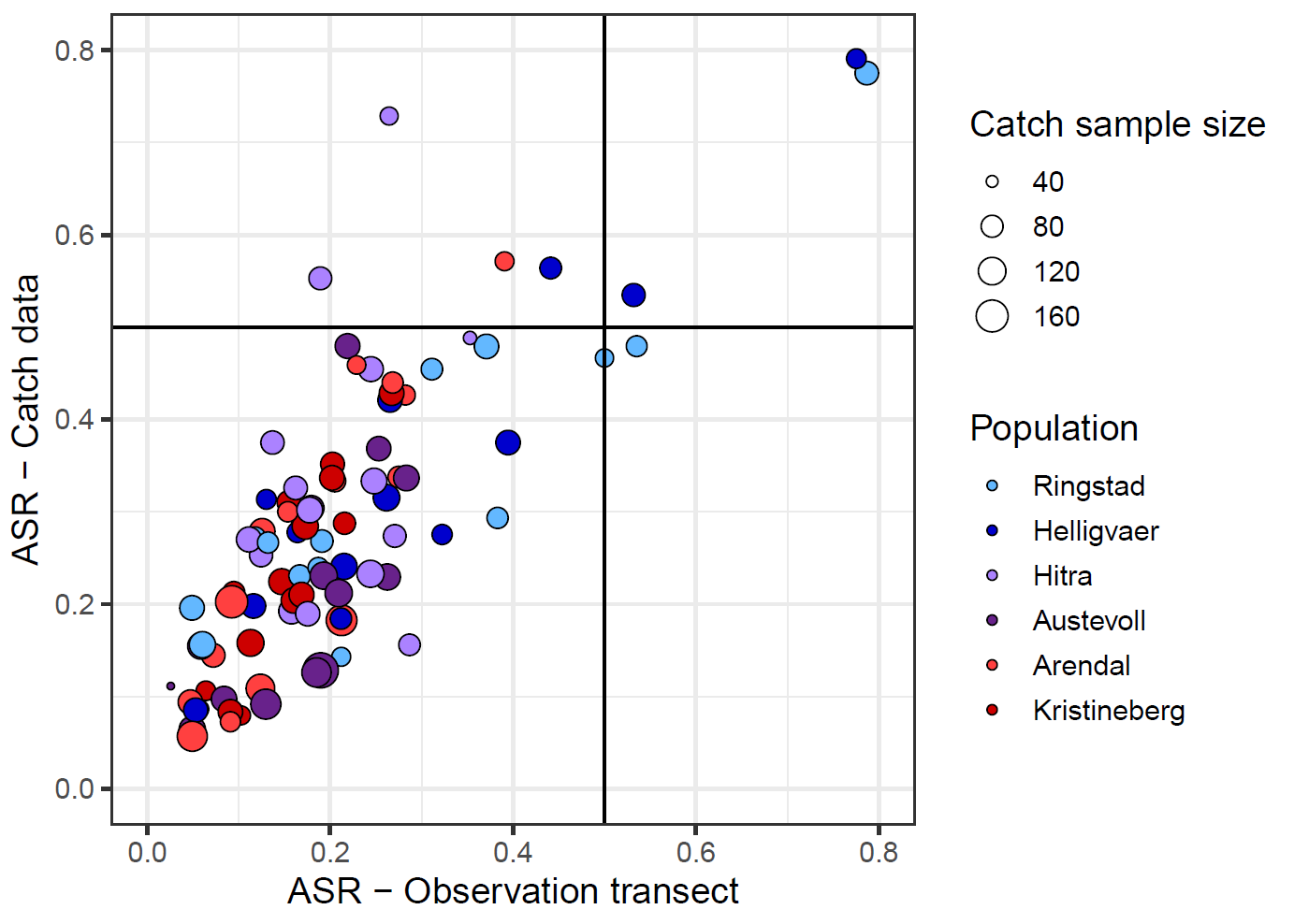

**Figure S13. Comparison ASR estimated from observation transects and catch data in the 2022 sampling effort.**

*The vertical and horizontal lines indicate an equal sex ratio. Each dot is a sampling occurrence where both an observation transect and a catch were conducted in sequence. Observation transects are done by a snorkelling observer, counting individuals of each sex. Catch consists in capturing individuals that are then brought to shore for phenotyping on land. Dot size represents catch sample size, and dot colour the population of sampling.*
